# Defined cellular reprogramming of androgen receptor-active prostate cancer to neuroendocrine prostate cancer

**DOI:** 10.1101/2025.02.12.637904

**Authors:** Shan Li, Kai Song, Huiyun Sun, Yong Tao, Arthur Huang, Vipul Bhatia, Brian Hanratty, Radhika A. Patel, Henry W. Long, Colm Morrissey, Michael C. Haffner, Peter S. Nelson, Thomas G. Graeber, John K. Lee

## Abstract

Neuroendocrine prostate cancer (NEPC) arises primarily through neuroendocrine transdifferentiation (NEtD) as an adaptive mechanism of therapeutic resistance. Models to define the functional effects of putative drivers of this process on androgen receptor (AR) signaling and NE cancer lineage programs are lacking. We adapted a genetically defined strategy from the field of cellular reprogramming to directly convert AR-active prostate cancer (ARPC) to AR-independent NEPC using candidate factors. We delineated critical roles of the pioneer factors ASCL1 and NeuroD1 in NEtD and uncovered their abilities to silence AR expression and signaling by remodeling chromatin at the somatically acquired AR enhancer and global AR binding sites with enhancer activity. We also elucidated the dynamic temporal changes in the transcriptomic and epigenomic landscapes of cells undergoing acute lineage conversion from ARPC to NEPC which should inform future therapeutic development. Further, we distinguished the activities of ASCL1 and NeuroD1 from the inactivation of RE-1 silencing transcription factor (REST), a master suppressor of a major neuronal gene program, in establishing a NEPC lineage state and in modulating the expression of genes associated with major histocompatibility complex class I (MHC I) antigen processing and presentation. These findings provide important, clinically relevant insights into the biological processes driving NEtD of prostate cancer.

## INTRODUCTION

Prostate cancer (PC) is a hormonally-driven disease in which androgen receptor (AR) signaling plays a fundamental role in defining a luminal epithelial cancer lineage and promoting cell survival and proliferation. Thus, androgen deprivation therapy (ADT) and AR signaling inhibitors (ARSIs) represent standard and effective frontline treatments for advanced PC. While the majority of PCs initially respond to combined treatment with ADT and an ARSI, therapeutic resistance ultimately develops over the course of months to years and leads to a disease state known as castration-resistant PC (CRPC) ^1,2^. The mechanisms by which CRPC subvert ADT and ARSIs are numerous, but most are centered on aberrant reactivation of AR signaling through AR gene mutations, amplification, alternative splicing, and enhancer duplication ^2–5^. Beyond reactivation of the targeted AR pathway, clinical evidence implicates lineage plasticity or a shift in cell identity as a more extreme adaptive mechanism of resistance in PC and other epithelial malignancies. A prime example is treatment-related transdifferentiation of AR-active PC (ARPC) to neuroendocrine PC (NEPC) in which the AR program is extinguished and supplanted with a NE program (NEtD) ^6,7^, thereby leading to a cancer lineage state that no longer depends on AR signaling and is resistant to ADT and ARSIs.

Multiple genetic and molecular events have been associated with the process of NEtD, including amplification of *MYCN*, loss of RE1-silencing transcription factor (REST) activity, and loss of *PTEN*, *TP53*, and *RB1* ^8–13^. While our prior studies using human prostate epithelial transformation models have demonstrated that overexpression of *MYCN* and constitutively active myristoylated *AKT1* ^8^ or the PARCB factors (dominant-negative *T<u>P</u>53*, myristoylated *<u>A</u>KT1*, short hairpin targeting *<u>R</u>B1*, *MY<u>C</u>*, and *<u>B</u>CL2*) can initiate NEPC ^9^, these experimental systems start with AR-null basal prostate epithelial cells. Thus, they do not address how AR-enforced luminal epithelial lineage commitment may be repressed and bypassed which is a critical step in the NEtD of PC. The leading biological model currently available to investigate NEtD is the LTL-331 ARPC patient-derived xenograft (PDX) model which reproducibly undergoes transdifferentiation after passage in castrated mice to the LTL-331R NEPC tumor model ^14^. Repeated biopsies and molecular analyses of tumors from this system have provided insights into molecular programs that are associated with NEtD ^15^. Yet, this model remains experimentally unwieldy for functional studies, and other more tractable approaches have been lacking.

Another challenge has been the inconsistent definition of NEPC applied in the field. Clinically, NEPC consists of a heterogeneous group of neuroendocrine tumors primarily defined by histologic morphology that includes a range of subtypes from well-differentiated carcinoid to aggressive large-cell and small-cell carcinomas ^16^. The expression of NE markers like chromogranin A (CHGA), synaptophysin (SYP), or CD56 (neural cell adhesion molecule 1 or NCAM1) are often assayed by immunohistochemistry (IHC) to support or confirm a diagnosis of NEPC. In general, individual NE markers expression can be heterogeneous, and combined marker scores are more robust ^17^. In experimental studies, phenotypic features such as NE marker expression or the emergence of dendrite-like projections from PC cells are broadly used as indicators of NE transdifferentiation of PC. The use of these sparse phenotypic features to define NEPC may be problematic as they may not reflect a *bona fide* cancer lineage state divergent from ARPC. A prime example is amphicrine prostate cancer (AMPC) which co-expresses AR and NE markers ^18^. Despite the expression of NE markers, AMPC models such as VCaP remain sensitive to the ARSI enzalutamide, indicating a functional dependence on the AR-driven lineage program.

Lineage-defining transcription factors and transcription factor networks specify cell states by regulating chromatin accessibility and gene expression programs. The basic helix-loop-helix (bHLH) proteins Achaete-scute homolog 1 (ASCL1) and neurogenic differentiation 1 (NeuroD1) are pioneer transcriptional factors that are necessary for lineage specification during neuronal development ^19,20^. In cancer, ASCL1 and NeuroD1 define distinct subtypes of small cell lung cancer (SCLC) ^21^ and NEPC ^22^ with differential transcriptional, epigenetic, cell surface proteomic, and drug sensitivity profiles ^23–25^. The role of ASCL1 as a driver of neuroendocrine transformation has been supported by the genetically engineered *TCKO* mouse model of high-grade neuroendocrine lung cancer where *Ascl1* knockout prevents tumor formation ^21^. Further, knockdown of *ASCL1* or *NEUROD1* in established human SCLC or NEPC cell line models significantly impairs cell viability ^26,27^, indicating that these factors are needed for sustained survival and growth. Insights into the functional role of these pioneer transcription factors in mediating the acute lineage conversion from ARPC to NEPC are limited. Recent work has shown that ASCL1 expression may be induced by ARSI therapy in PC and ASCL1 activates a neuronal stem cell-like lineage program through chromatin remodeling mediated by the polycomb repressive complex 2 (PRC2) ^28^.

In this study, we developed a genetically defined strategy adapted from the field of cellular reprogramming to directly convert ARPC to NEPC. We identified key functional roles for ASCL1 and NeuroD1 in NE transdifferentiation, specifically new findings uncovering their ability to silence the AR program through chromatin remodeling of the AR enhancer which is somatically amplified in PC and global AR binding sites with enhancer activity. In contrast, we found that loss of REST activity induces the expression of NE markers but is insufficient to drive a NE cancer lineage state. Repeated sampling over the reprogramming time course revealed a wide range of temporal changes in the transcriptomic and epigenomic landscapes associated with the lineage conversion from ARPC to NEPC. Lastly, we found that ASCL1 and NeuroD1 expression, but not the loss of REST activity, induces the downregulation of class I major histocompatibility complex (MHC) expression in PC, providing a basis for the lineage-specific potential for immune evasion associated with NEPC.

## RESULTS

### Candidate factors directly reprogram ARPC to NEPC and bypass the dependence on AR signaling

To functionally define factors required for the conversion of ARPC to NEPC, we established a NEtD assay wherein candidate factors were introduced into the human ARPC cell lines LNCaP and C4-2B after which they were propagated in media conditions permissive to the growth of NEPC and analyzed for a set of phenotypic markers of ARPC and NEPC after two weeks (**Figure 1A**). Initially, we introduced a pool of lentiviruses (LVs) expressing eight factors associated with NEPC including dominant-negative *TP53* R175H, a short hairpin targeting *RB1* (shRB1), *MYCN*, *ASCL1*, *SRRM4*, *NR0B2*, *BCL2*, and *KRAS* G12V. Loss of *TP53* and *RB1* as well as amplification of *MYCN* have been identified as common genetic features associated with NEPC ^8,9,29,30^. SRRM4 is a splicing regulator that has been shown to inactivate the RE-1 silencing transcription factor (REST) which functions to suppress a major neural gene program. Loss of REST has been implicated as a common feature in NEPC that has been reported to functionally drive NEtD of PC ^12,31^. In addition, the pioneer neural transcription factor ASCL1 is expressed in most NEPC and appears to define a transcriptional and epigenetic subtype of the disease ^22^. NR0B2, otherwise known as short heterodimer partner (SHP), is an orphan nuclear receptor that is highly expressed in NEPC that interacts with AR and inhibits AR activity ^32^. Lastly, BCL2 and KRAS G12V were included to potentially reduce apoptosis and enhance reprogramming efficiencies ^33,34^.

**Figure 1.**
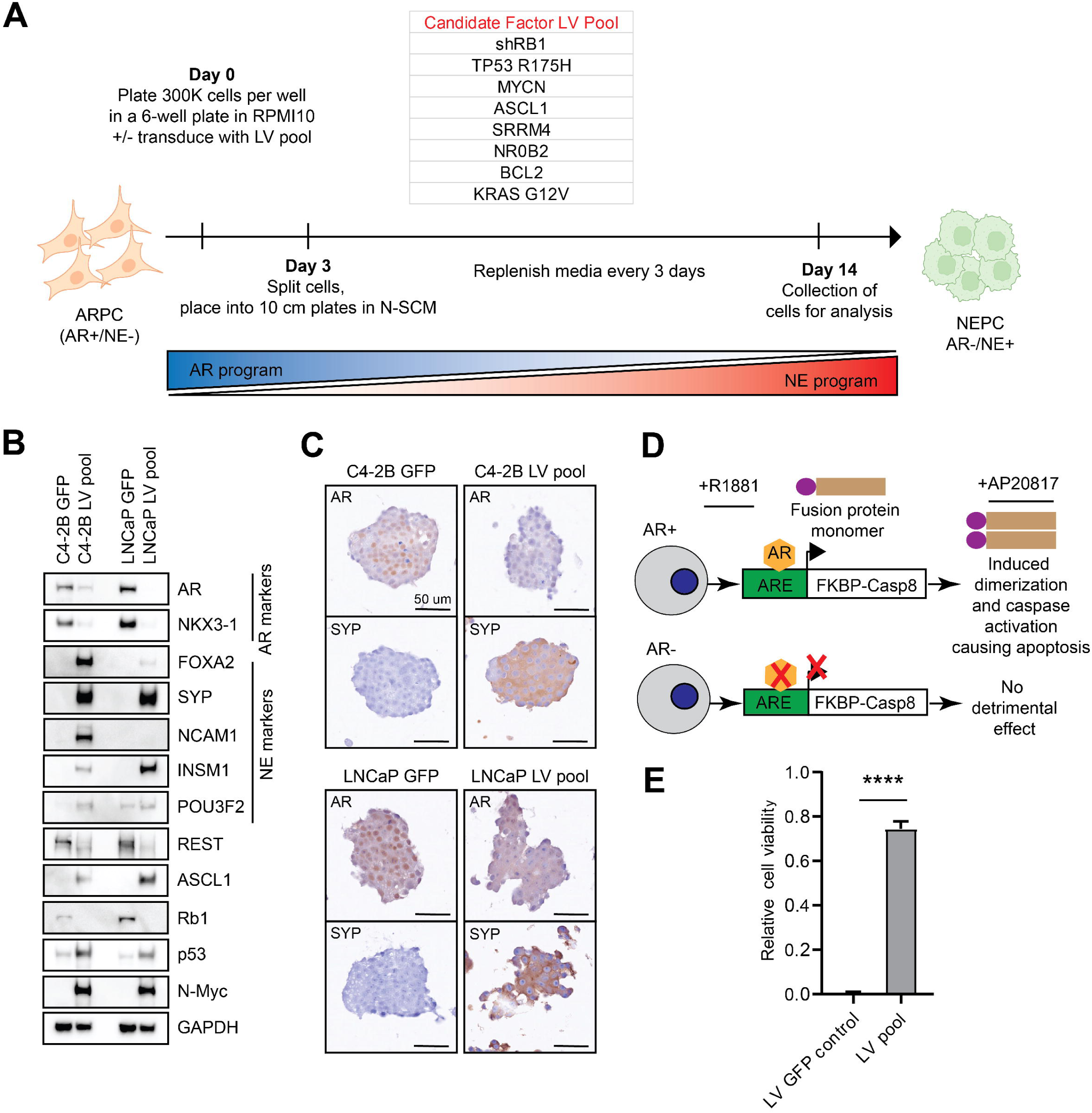
Direct reprogramming of ARPC to NEPC with functional bypass of a dependence on AR signaling. (**A**) Experimental schema of the reprogramming of ARPC cells to NEPC through the introduction of a candidate factor lentiviral (LV) pool and propagation in neural stem cell media (N-SCM) permissive to NEPC growth. (**B**) Immunoblot analysis of the C4-2B and LNCaP cell lines transduced with either green fluorescent protein (GFP) or the LV pool and subjected to the reprogramming assay. Lysates were collected on day 14. (**C**) Immunocytochemical analysis of the C4-2B and LNCaP lines at day 14. (**D**) Experimental schema for the stringent negative selection of AR^+^ cell populations based on the AR-dependent expression of an inducible FKBP-Casp8 fusion protein that dimerizes and induces caspase activation and apoptosis in the presence of the FK506 analog AP20817. (**E**) Relative cell viability determined by CellTiter-Glo assay of treated versus untreated groups after five days (n=4 each). P-value was assessed by Student’s t-test. **** denotes p <0.0001.

The introduction of this candidate factor LV pool at a multiplicity-of-infection (MOI) of four for each factor in this assay led to a significant downregulation of AR and the AR target NK3 homeobox 1 (NKX3-1) as well as the upregulation of NE markers including forkhead box A2 (FOXA2), SYP, NCAM1, insulinoma-associated protein 1 (INSM1), and POU class 3 homeobox 2 (POU3F2 or BRN2) by immunoblot analysis (**Figure 1B**). Furthermore, immunocytochemical analysis of the C4-2B and LNCaP cell lines transduced with the candidate factor LV pool demonstrated a prominent loss of nuclear AR expression and increased SYP expression (**Figure 1C**). The AR^low/-^NE^+^ cancer phenotype induced by the candidate factor LV pool was durable based on IHC studies in tumors obtained after subcutaneous xenografting in non-castrate male NOD.Cg-*Prkdc^scid^ II2rg^tm1Wjl^ SzJ* (NSG) mice (**Figure S1A**).

We next sought to determine whether the PC cells reprogrammed with the candidate factor LV pool were functionally independent of AR signaling. We implemented a potent AR-dependent negative selection strategy (**Figure 1D**) wherein C4-2B cells were transduced with LV encoding AR response elements (ARE) upstream of a FK506 binding protein-caspase 8 (FKBP-Casp8) fusion. A clonal C4-2B ARE-FKBP-Casp8 line was established and transduced with either a control LV expressing green fluorescent protein (GFP) or the candidate factor LV pool and subjected to selection with 1 nM R1881 (to induce AR signaling and expression of FKBP-Casp8 in AR^+^ cells) and AP20187 which is a synthetic FK506 analog (to activate FKBP-Casp8 and induce apoptosis). We identified no viable, adherent cells in the control LV condition after five days of selection. In contrast, we appreciated viable, adherent cells in the candidate factor LV pool condition and ∼70% of cells demonstrated functional AR bypass as they could survive stringent selection against AR signaling (**Figures 1E and S1B**). These finding indicated that a discrete set of factors is capable of reprogramming human ARPC, in the context of negative selection for non-reprogrammed cells (modeling highly stringent AR inhibitor therapy), to a cellular state that recapitulates critical phenotypic and functional features of NEPC.

### The pioneer neural transcription factor ASCL1 suppresses the AR program during NEtD but requires additional genetic alterations to maintain a proliferative phenotype

To specify the functional contribution of each candidate factor in the LV pool to NE reprogramming, we employed a leave-one-out approach in which each factor was iteratively left out from the LV pool in the C4-2B (**Figure 2A**) and LNCaP **(Figure S2)** cell lines. Based on immunoblot analyses, we observed several established regulatory interactions that have been previously identified in cancer. These included 1) ectopic expression of N-Myc leading to elevated levels of NCAM1 in neuroblastoma ^35^, 2) repression of POU3F2 expression by Rb1 in retinoblastoma ^36^, 3) and enhanced SOX2 expression with Rb1 inactivation in PC ^13^. Most strikingly, we observed that the exclusion of ASCL1 diminished the decrease in AR and NKX3-1 expression and likewise abrogated the otherwise induced FOXA2, NCAM1, and INSM1 expression, implicating ASCL1 as a critical player in NE transdifferentiation. In contrast, the omission of SRRM4 restored REST expression and suppressed SYP but had little effect on AR or other NE markers. These data are consistent with prior reports in which ectopic introduction of SRRM4 in PC cell lines induced SYP and chromogranin B expression but did not directly affect AR expression ^31,37^. Removal of NR0B2, BCL2, and KRAS G12V had no significant effects on AR and NE marker expression (**Figures 2A and S2A**).

**Figure 2.**
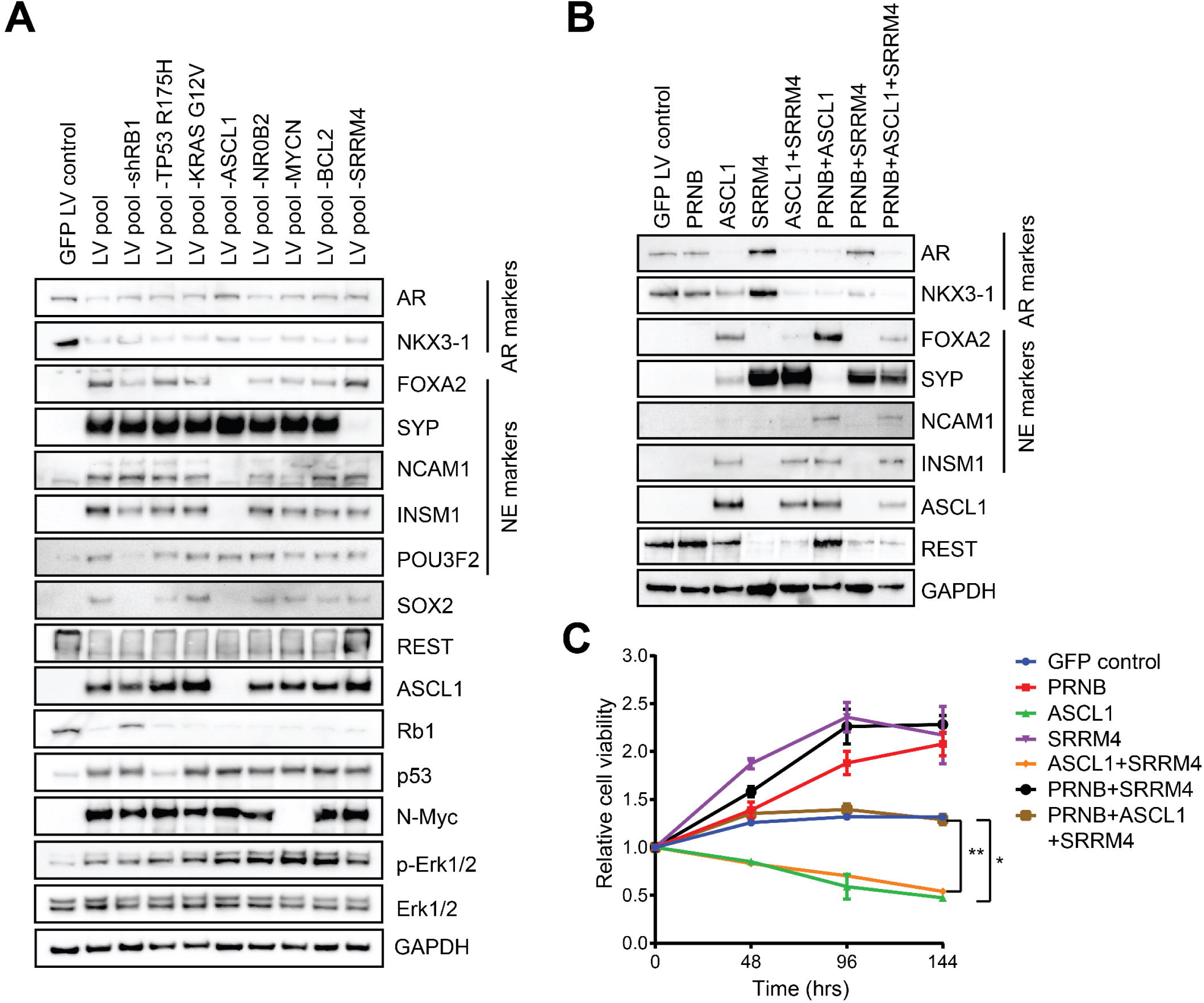
The pioneer neural transcription factor ASCL1 suppresses AR expression and drives NE transdifferentiation in prostate cancer. Immunoblot analysis of (**A**) leave-one-out conditions and (**B**) factor reconstitution conditions in reprogramming studies using the C4-2B cell line. PRNB represents dominant-negative T<u>P</u>53 H175R, sh<u>R</u>B1, MYC<u>N</u>, and <u>B</u>CL2. Lysates were collected on day 14. (**C**) Relative cell viability over time determined by CellTiter-Glo assay of C4-2B cells reprogrammed with various factor combinations (n=4 per condition). * denotes p <0.05, ** denotes p <0.01.

Next, we employed an orthogonal factor reconstitution strategy to identify the minimal set of factors needed to induce NEtD in the ARPC lines. Immunoblot analyses showed that ASCL1 expression alone downregulated AR markers and induced NE marker expression **(Figure 2B)**. On the other hand, the PRNB factors (dominant-negative T<u>P</u>53 H175R, sh<u>R</u>B1, MYC<u>N</u>, and <u>B</u>CL2) were insufficient to enforce NEtD (**Figure 2B**). SRRM4 alone imparted a partial effect on phenotype as it led to a loss of REST expression and upregulated SYP without reducing AR markers (**Figure 2B)**. As *TP53*, *RB1*, *MYCN*, and *BCL2* alterations are common in NEPC and other aggressive NE carcinomas we questioned why they appeared to be dispensable in our NEtD assay. To confirm whether all factors were introduced into the cells, we encoded each lentiviral construct with a unique 10-nucleotide barcode and performed single-cell DNA amplicon sequencing ^38^ 72 hours after transduction. Of 3,870 cells analyzed, 3,785 cells (97.8%) demonstrated reads from all barcodes, while 84 cells (2.2%) were missing one barcode and 1 cell was missing two barcodes (**Figure S2B**). We measured cell proliferation for each of the factor reconstitution conditions as an indicator of proliferative fitness in our experimental system. PRNB significantly enhanced proliferation while ASCL1 was detrimental to cell viability over time in the ARPC lines (**Figure 2C and Figure S2C**). This effect of ASCL1 is consistent with prior reports that ASCL1 promotes cell cycle exit and terminal differentiation when overexpressed in neural progenitor cells and glioblastoma stem cells ^39^. Notably, SRRM4 promoted proliferation but alone was insufficient to rescue the effects of ASCL1 without PRNB. These data support the concept that the appropriate genetic and proliferative context may be an important determinant of ASCL1’s impact on the emergence of NEPC and highlight the idea that overall fitness may be a combination of proliferation- and lineage-linked benefit. In other words, this mechanism of therapeutic escape may require a change in lineage without a substantial decrease in proliferative fitness.

### NeuroD1 is competent to induce neuroendocrine lineage reprogramming

We have previously observed that most NEPCs can be assigned to ASCL1^high^, NeuroD1^high^, or mixed ASCL1/NeuroD1 groups based on transcriptional profiling ^26^. These findings have also been described by other groups wherein ASCL1 and NeuroD1 appear to define molecular subtypes of NEPC ^22^ and the neuroendocrine subtypes of small cell lung cancer ^21^. To determine whether NeuroD1 can also drive lineage conversion from ARPC to NEPC, we performed the reprogramming assay using the C4-2B and LNCaP ARPC lines by replacing ASCL1 with NeuroD1 or combining NeuroD1 with ASCL1. The addition of NeuroD1 to PRNB and SRRM4 substantially diminished AR markers and increased NE marker expression (**Figure 3A**). We discovered that NeuroD1 induced ASCL1 expression and heightened overall NE marker panel expression. Bulk RNA-seq gene expression analysis of the reprogrammed cell lines with respect to established 22-gene AR signature and NE signature scores revealed a significant de-enrichment of the AR gene signature and enrichment of the NE gene signature in the PRNBSA (<u>PRNB</u>, <u>S</u>RRM4, and <u>A</u>SCL1) and PRNBSN (<u>PRNB</u>, <u>S</u>RRM4, and <u>N</u>euroD1) conditions relative to the GFP control (**Figure 3B**). As C4-2B is derived from the LNCaP cell line, we also confirmed the reproducibility of these findings in the independent ARPC cell line MDA PCa 2b using this cellular reprogramming assay (**Figure S3A, C**).

**Figure 3.**
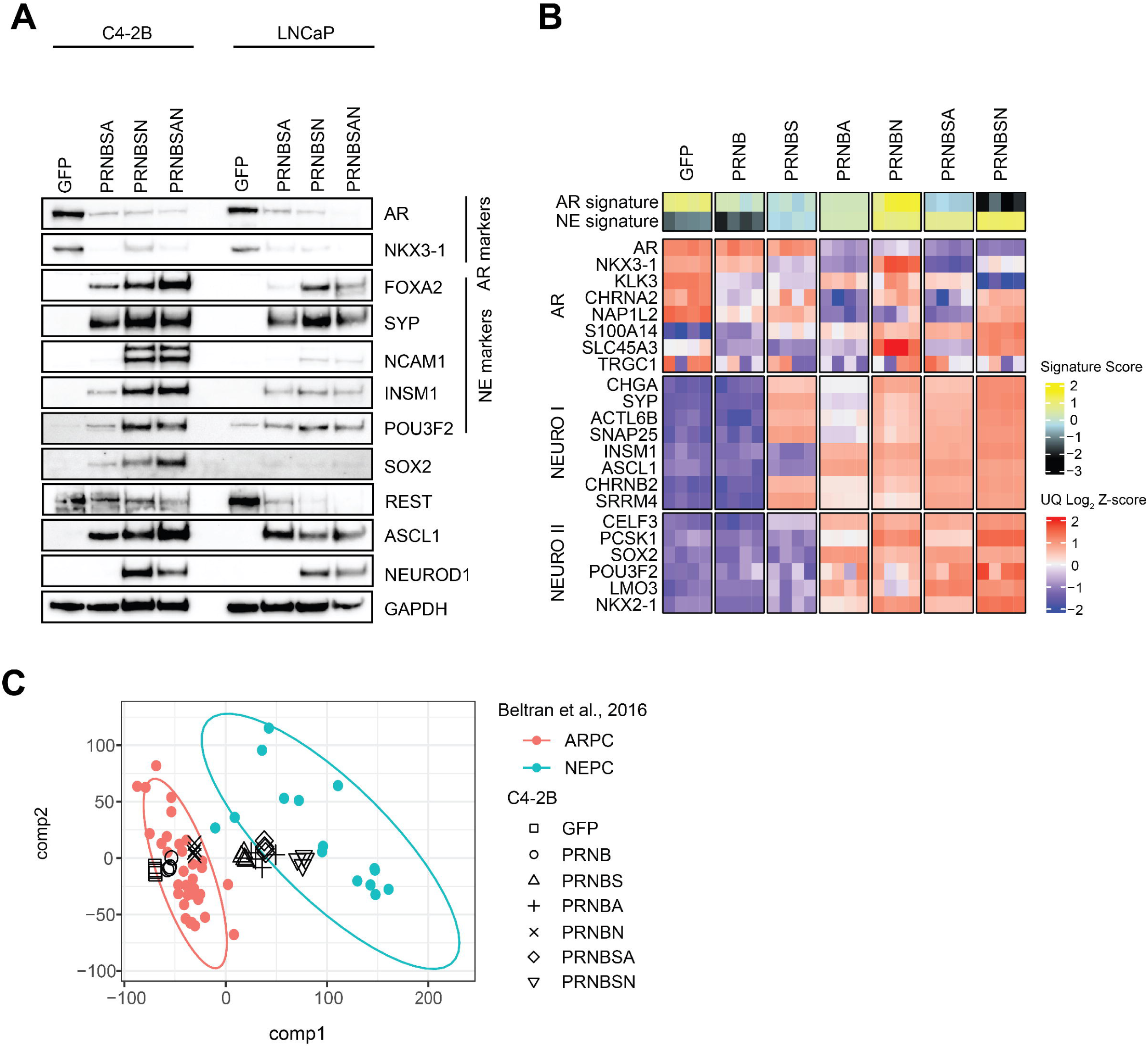
ASCL1 and NeuroD1 are competent to induce NE lineage reprogramming of prostate cancer. **(A)** Immunoblot analysis of C4-2B and LNCaP cells reprogrammed with GFP or PRNB, SRRM4, and ASCL1 and/or NeuroD1. Lysates were collected on day 14. (**B**) Heatmap of RNA-seq gene expression from reprogrammed C4-2B cell line conditions showing 22-gene AR and NE signature scores and select genes associated with the AR program and NE programs (NEURO I and NEURO II). UQ: upper quartile normalization. (**C**) Partial least squares-discriminant analysis (PLS-DA) plot based on RNA-seq gene expression of reprogrammed C4-2B cell line conditions (black shapes) projected onto human ARPC (red dots) and NEPC (blue dots) samples from Beltran et al., 2016. Ellipses represent 95% confidence level for multivariate t-distributions defined by ARPC (red) and NEPC (blue) data.

Several neural transcription factors including FOXA2, INSM1, POU3F2, and SOX2 have also been described as documented drivers or regulators of NE transdifferentiation in PC and other cancer types ^13,40,41^. We sought to determine whether these factors are independently competent to induce NE lineage reprogramming. We introduced PRNB, SRRM4, and each of the above listed neural transcription factors individually to the C4-2B line in the NEtD assay. In contrast to ASCL1 and NeuroD1, these factors were by themselves incapable of NE lineage reprogramming (**Figure S3B**). As ASCL1 and NeuroD1 also appear to regulate the expression of FOXA2 and INSM1, ASCL1 and NeuroD1 may sit atop a hierarchy of multiple NE-associated transcription factors required in combination to establish and maintain a NEPC transcriptional regulatory network.

To confirm the relevance of our NE transdifferentiation models in PC and to use omic-scale data to minimize the effect of individual marker heterogeneity, we performed partial least squares-discriminant analysis (PLS-DA) of RNA-seq gene expression data from previously characterized human metastatic CRPC patient specimens annotated as ARPC or NEPC ^42^ and projected the C4-2B factor-reconstituted samples onto them (**Figure 3C**). This analysis showed that PRNB alone did not induce a substantial shift away from the ARPC phenotype. Further, the addition of PRNB and SRRM4 (PRNBS) resulted in a minor, intermediate shift toward the NEPC phenotype. Only upon the addition of ASCL1 or NeuroD1 to PRNB (PRNBA and PRNBN) or PRNBS (PRNBSA and PRNBSN) did we observe a dramatic gene expression program-based shift from ARPC to NEPC. Similar results were observed in in MDA PCa 2b cell line (**Figure S3D**). These data highlight the relative lineage impacting importance of these factors in NEtD of PC.

SRRM4 promotes alternative splicing and inactivates REST through the inclusion of a neural exon that generates the REST-4 isoform ^31,43^. We confirmed the presence of alternatively spliced *REST* mRNA in C4-2B reprogramming conditions that included SRRM4 (**Figure S4A**). Further, we performed PLS-DA analysis on alternative splicing data from Beltran et al., 2016 and projected the C4-2B reprogrammed samples onto them. This analysis indicated that SRRM4 drives the alternative splicing phenotype associated with NEPC with a marginal contribution from ASCL1 or NeuroD1 (**Figure S4B**).

### Dynamic changes in cancer phenotype, transcriptional and epigenetic landscapes during NE reprogramming

Having established a defined and tractable system that recapitulates NE transdifferentiation of PC, we sought to investigate the dynamic molecular and regulatory changes underlying this shift in cancer lineage state. We applied the NE reprogramming assay to the ARPC C4-2B line and temporally sampled multiple timepoints within the 14-day reprogramming period (D2, D3, D4, D5, D8, D11, and D14) for downstream analyses (**Figure 4A**). Immunoblot analyses revealed a reduced expression of AR markers (AR and NKX3-1) as early as D2, with expression of the NE markers INSM1 and FOXA2 evident shortly thereafter on D4-5 (**Figure 4B**). These findings indicate that downregulation of the AR program precedes the establishment of the NE program in our experimental system. Transcriptome profiling by bulk RNA-seq of the C4-2B cells reprogrammed with PRNBSA or PRNBSN recapitulated the findings from the immunoblot analyses where downregulation of *AR* expression preceded the induction of *INSM1* expression (**Figures 4C and D**). Notably, PCA analysis of the time course RNA-seq gene expression data from C4-2B cells reprogrammed with PRNBSA or PRNBSN, but not PRNBS, showed an arc-like trajectory during NEtD, indicating dynamic changes in gene expression programs during ARPC to NEPC reprogramming (**Figures S5A-C**). This arc-like trajectory is also broadly observed in normal and cancer development and cellular differentiation ^44–47^.

**Figure 4.**
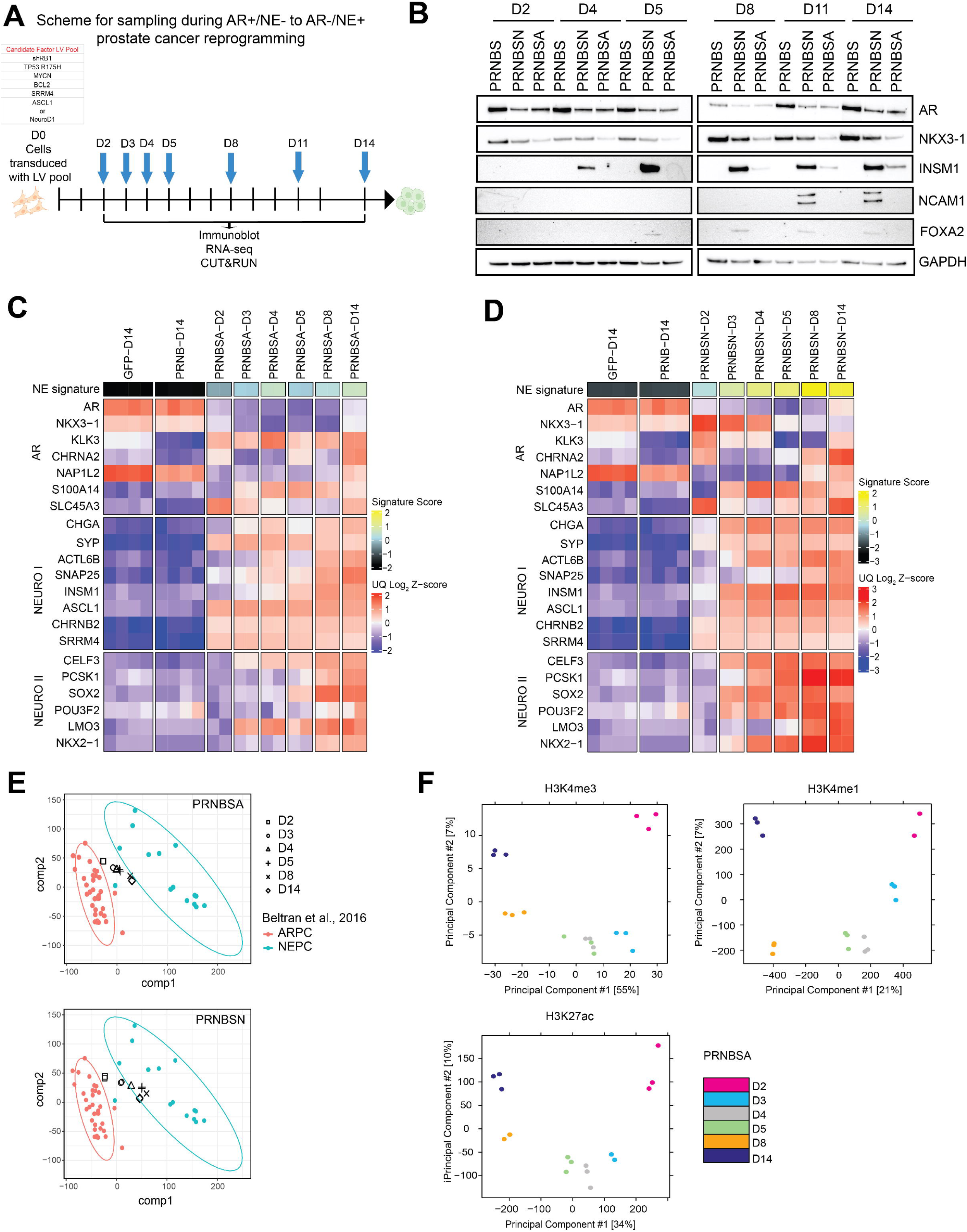
Dynamic changes in cancer phenotype and transcriptional and epigenetic landscapes during acute reprogramming of ARPC to NEPC. (**A**) Experimental schema for the temporal investigation of the reprogramming of ARPC to NEPC by immunoblot, RNA-seq, and CUT&RUN analyses. (**B**) Immunoblot analysis of C4-2B cells modified with PRNB and SRRM4 (PRNBS), PRNBS and NeuroD1 (PRNBSN), or PRNBS and ASCL1 (PRNBSA) that were collected at various timepoints during the 14-day reprogramming period. Heatmaps of RNA-seq gene expression from C4-2B cells reprogrammed with (**C**) PRNBSA or (**D**) PRNBSN over the 14-day reprogramming period showing 22-gene NE signature scores and select genes associated with the AR program and NE programs (NEURO I and NEURO II). (**E**) PLS-DA plot based on RNA-seq gene expression of C4-2B cells reprogrammed with PRNBSA (top) or PRNBSN (bottom) at different timepoints (black shapes) projected onto human ARPC (red dots) and NEPC (blue dots) samples from Beltran et al., 2016. Ellipses represent 95% confidence of t-distribution defined by ARPC (red) and NEPC (blue) data. (**F**) Principal component analysis (PCA) of CUT&RUN enrichment for H3K4me3, H3K4me1 and H3K27Ac signal in C4-2B cells reprogrammed with PRNBSA over time.

We next used PLS-DA projections from the time course RNA-seq gene expression data to further investigate the temporal nature of the phenotypic shift from ARPC to NEPC during NEtD. We distinguished three general phases of reprogramming in the C4-2B cells modified with PRNBSA and PRNBSN that could be defined as “early” on D2, “intermediate” from D3-5, and “late” from D8-14 (**Figure 4E**). In contrast, these phases were not apparent in C4-2B cells transduced with PRNB and SRRM4 (**Figure S5D**), further supporting the previous finding that ASCL1 or NEUROD1 is needed for full lineage shift via NEtD.

Lineage reprogramming of ARPC to NEPC is largely driven by epigenetic dysregulation and reprogramming as few genetic alterations apart from the enrichment of *TP53* and *RB1* mutations and genomic amplification of *MYCN* have been identified in NEPC ^30,42,48,49^. To investigate global changes in the epigenetic landscape during the transition from ARPC to NEPC, we performed Cleavage Under Targets & Release Using Nuclease (CUT&RUN) analysis on PRNBSA reprogrammed C4-2B cells over the 14-day time course to examine genome-wide changes in histone modifications, including monomethylation of histone H3 at lysine 4 (H3K4me1) which is commonly associated with gene enhancers, acetylation of histone H3 on lysine 27 (H3K27ac) which indicates both active promoters and enhancers ^50,51^, and trimethylation of histone H3 at lysine 4 (H3K4me3) which marks active gene promoters. Unsupervised principal component analysis (PCA) showed that these histone modifications shift significantly during the time course (**Figure 4F**), indicating that dynamic epigenetic changes occur during NEtD.

As ASCL1 is a pioneer transcription factor, we queried whether there may be time-dependent changes in putative ASCL1-mediated gene regulation during the acute conversion of ARPC to NEPC. We integrated the RNA-seq gene expression data and ASCL1 CUT&RUN genome occupancy data over the 14-day reprogramming period to identify genes both in proximity to ASCL1 binding and upregulated in PRNBSA compared to PRNBS. Gene set enrichment analysis (GSEA) revealed several pathways enriched early during NE reprogramming including the CDC25 pathway, TGFβ1 pathway, and estradiol response pathway (**Figure 5A**). In contrast, ASCL1 targets, PRC2 targets including those of the core subunits embryonic ectoderm development (EED) and SUZ12, and neuronal fate commitment pathways were significantly enriched later in the NE transdifferentiation process. The pathways enriched early may represent components of an unstable transcriptional state prior to the establishment of a stable NEPC lineage, consistent with our observation of an arc-like trajectory.

**Figure 5.**
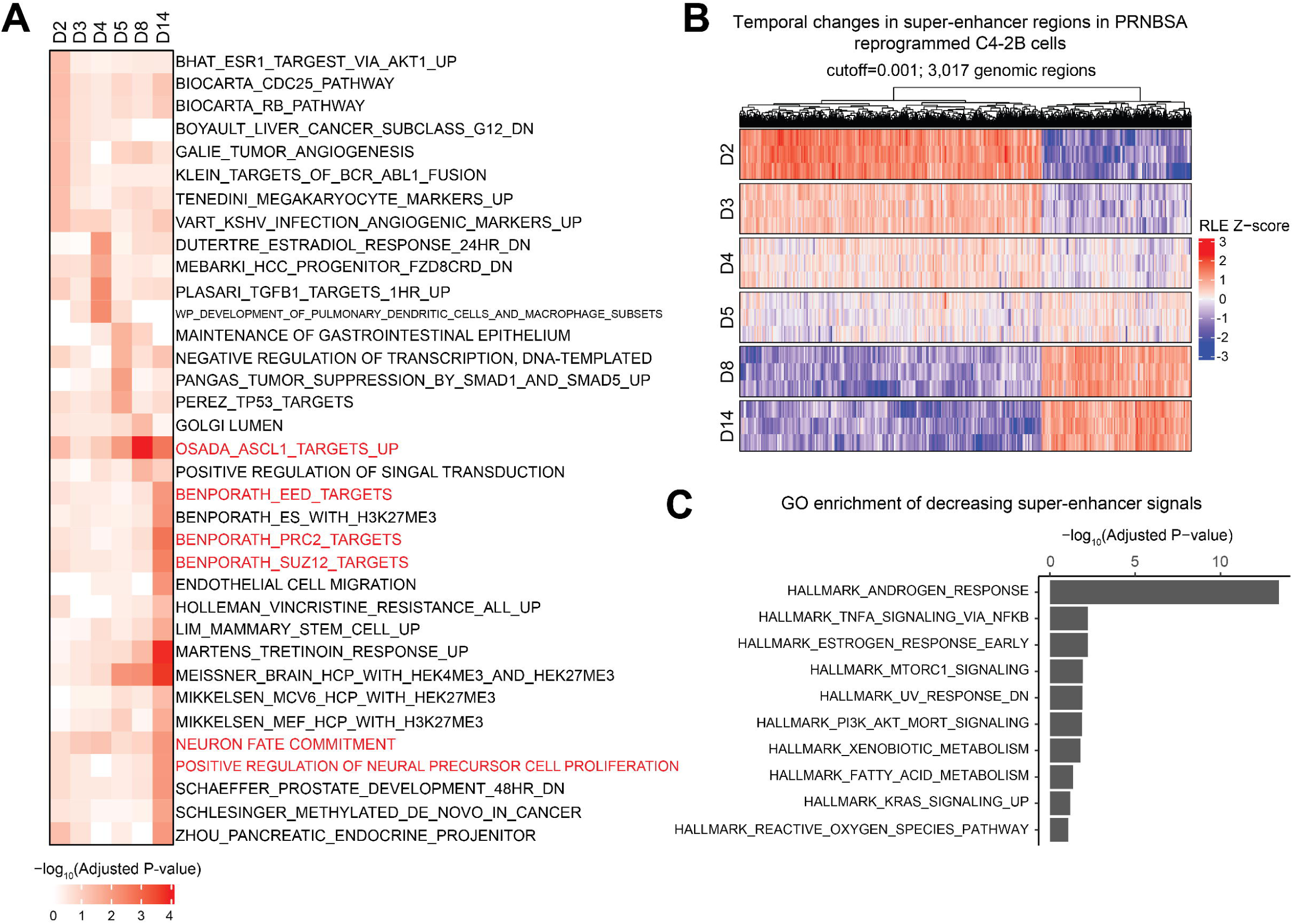
Time-dependent changes in ASCL1-regulated gene programs and super-enhancer organization during NE transdifferentiation of prostate cancer. **(A)** Heatmap of MSigDB gene sets enriched over time with ASCL1-regulated genes identified by integrating CUT&RUN and RNA-seq data from C4-2B cells reprogrammed with PRNBSA over the 14-day reprogramming period. (**B**) Heatmap showing dynamic changes in super-enhancer regions in C4-2B cells reprogrammed with PRNBSA over the 14-day reprogramming period. Genomic regions shown were obtained by setting a p-value cutoff of 1e-03 on Kendall correlation between day and peak activity. RLE: Relative Log Expression normalization. (**C**) Plot showing Gene Ontology (GO) enrichment of Hallmark gene sets associated with decreasing super enhancer signals from D2 to D14 in C4-2B cells reprogrammed with PRNBSA.

Next, we evaluated the dynamic activities of active enhancers, super-enhancers, and active promoters in the NE reprogramming assay. We defined active enhancers based on overlapping H3K4me1 and H3K27ac peaks and characterized super-enhancers by applying the Rank Ordering of Super-Enhancer (ROSE) algorithm to H3K27ac peaks over the reprogramming time course ^52^. This analysis revealed a substantial temporal shift in active enhancers (**Figure S6A**) and super-enhancers (**Figure 5B**). Gene ontology (GO) analysis of downregulated active enhancer- and super-enhancer-associated genes over time revealed a highly significant enrichment of androgen response genes including the Hallmark Androgen Response gene set (**Figures S6B and 5C**). In contrast, genes associated with upregulated active enhancers over time were enriched for endocrine therapy resistance pathways and targets of the PRC2 complex core subunits SUZ12 and enhancer of zeste homolog 2 (EZH2) (**Figure S6C**). We also examined active promoters by determining the overlap of H3K4me3 and H3K27ac peaks over the reprogramming time course (**Figure S7A**). Genes associated with downregulated active promoters over time were enriched in androgen response and AR regulated targets (**Figure S7B**). On the other hand, GO analysis of upregulated active promoters over time did not highlight any gene sets of statistical significance.

### ASCL1/NeuroD1 inhibits AR expression by remodeling the somatically acquired AR enhancer

To further investigate how ASCL1 and NeuroD1 may suppress AR expression and signaling during NE reprogramming, we performed single-cell (sc) multiome RNA-seq and ATAC-seq analysis on D14 after reprogramming of C4-2B cells with PRNB, PRNBS, PRNBSA, and PRNBSN. The PRNB and PRNBS conditions were included as controls as these factor combinations have no apparent effect on the AR program. Pseudotime analysis revealed a progression from PRNB to PRNBS to PRNBSA to PRNBSN which was associated with a heightened NEPC score and reduced *AR* expression (**Figure S8A-D**). In addition, we observed two distinct clusters within the PRNBSA and PRNBSN conditions indicating heterogeneous cell populations likely due to differences in expression levels of *ASCL1* and *NEUROD1* (**Figures S8E, F**).

Most strikingly, the integration analysis of RNA-seq and ATAC-seq data identified a significant reduction in chromatin accessibility of the somatically acquired *AR* enhancer, but not the *AR* gene body, that was associated with reduced *AR* expression in the PRNBSA and PRNBSN conditions (**Figure 6A**). In contrast, the reduction in chromatin accessibility of the somatically acquired *AR* enhancer was not seen in the control PRNB and PRNBS conditions (**Figure 6A**). The somatically acquired enhancer of *AR* and its amplification independent of the *AR* gene body has previously been shown to be a noncoding driver of CRPC ^5^. Attesting to the clinical relevance of this finding, we identified a hepatic metastasis from a patient with lethal mCRPC who underwent rapid autopsy as part of the University of Washington Tissue Acquisition Necropsy (UW TAN) program that showed a mixed population of AR^+^/ASCL1^-^ and AR^-^/ASCL1^+^ tumors cells by immunofluorescence staining (**Figure 6B**). Similar to the findings from our NE reprogramming studies, scATAC-seq analysis of this metastatic tumor showed loss of chromatin accessibility at the enhancer of *AR* but not the *AR* gene body in AR^-^/ASCL1^+^ tumor cells compared to AR^+^/ASCL1^-^ cells (**Figure 6C**).

**Figure 6.**
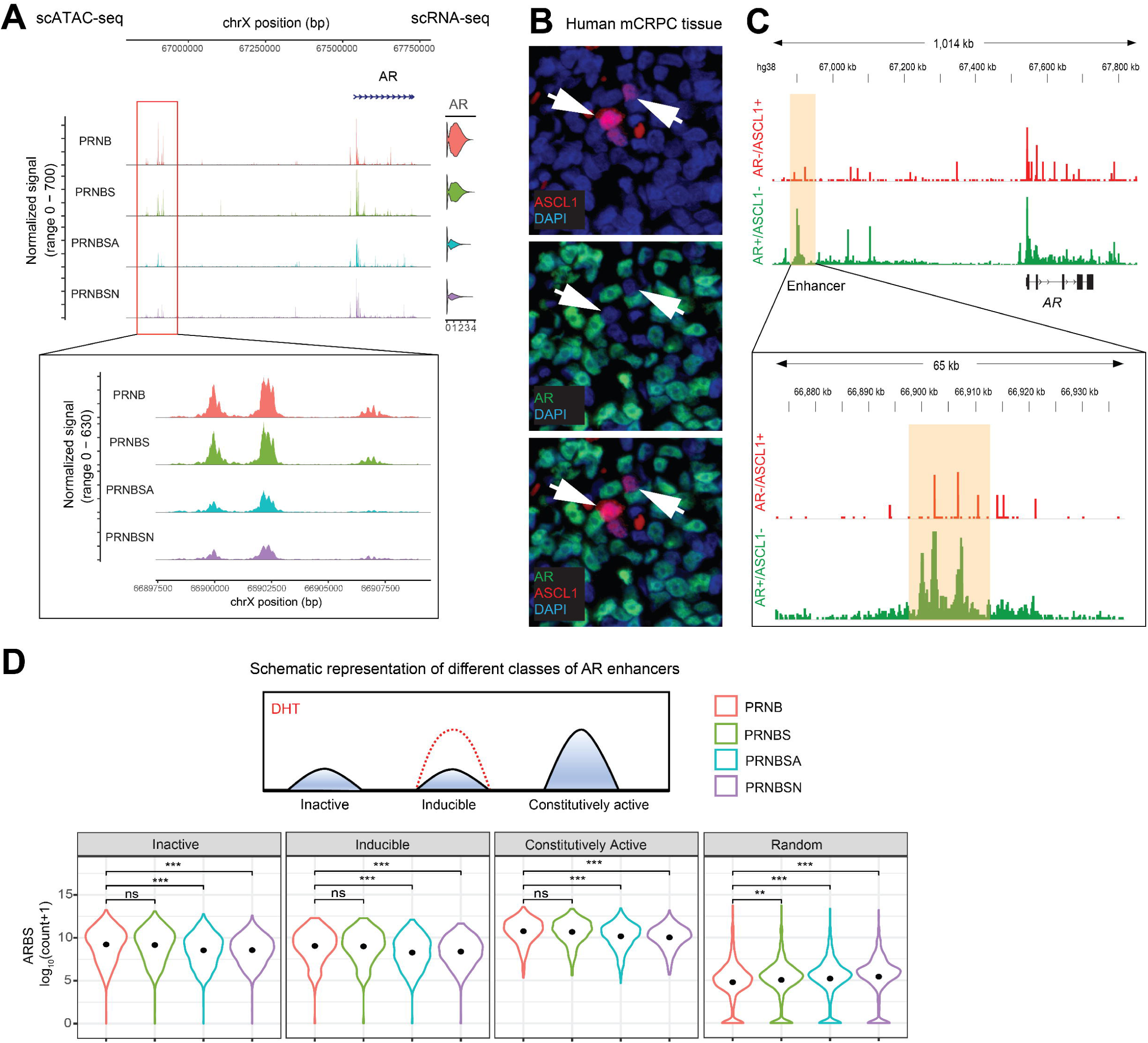
ASCL1/NeuroD1 inhibit AR expression by remodeling chromatin accessibility at the somatically acquired AR enhancer and global AR binding sites with enhancer activity. (**A**) Tracks showing chromatin accessibility peaks at the AR locus and AR gene expression (violin plot) from pseudo-bulk analysis of scATAC-seq and scRNA-seq data from C4-2B cells reprogrammed with PRNB, PRNBS, PRNBSA, and PRNBSN. Tracks in the region of the somatically acquired AR enhancer are magnified. (**B**) Immunofluorescence analysis of a human prostate cancer hepatic metastasis with ASCL1 staining (top), AR staining (middle), and an overlay of ASCL1 and AR staining (bottom) shown. DAPI was used as a nuclear counterstain. (**C**) Tracks showing chromatin accessibility at the AR locus by pseudo-bulk analysis of scATAC-seq data from the AR^-^/ASCL1^+^ and AR^+^/ASCL1^-^ tumor cell populations present in the human prostate cancer hepatic metastasis shown in **B**. Tracks in the region of the somatically acquired AR enhancer are magnified. (**D**) Schematic representation of different classes of AR enhancers (top) is shown. The distribution of chromatin accessibility at AR bindings sites (ARBS) of different enhancer activity and at random genomic regions of the same interval are shown. Black dots represent the means. ns denotes p >0.05, ** denotes p <0.01, *** denotes p <0.001.

We next sought to investigate the chromatin accessibility of genome-wide AR binding sites (ARBS) with enhancer activity. ARBS with enhancer activity have previously been characterized using Self-Transcribing Active Regulatory Regions sequencing (STARR-seq) in the LNCaP cell line and classified as constitutively active enhancers, inducible enhancers, and inactive enhancers based on their responsiveness to the androgen dihydrotestosterone ^53^. We evaluated the chromatin accessibility at these ARBS and identified a global reduction in the accessibility of ARBS with enhancer activity from least reprogrammed (PRNB) to most reprogrammed (PRNBSA or PRNBSN) cell lines (**Figure 6D**). In contrast, a random selection of genomic sites of similar intervals across all chromosomes, as a control, did not show reduction in chromatin accessibility. These results further demonstrate the acute epigenetic rewiring of PC during NEtD to silence an AR-driven PC lineage program.

### ASCL1/NeuroD1 but not loss of REST via SRRM4 activity establish a NE transcriptional network

Lineage-defining transcription factors including neural lineage transcriptions factors have been reported to auto-activate their own expression by binding to super-enhancers to establish a positive feedback loop required to sustain a lineage program ^54^. We investigated the potential for self-regulation of ASCL1 and NeuroD1 in our NE reprogramming assay based on differential RNA-seq read alignment of exogenous proviral transcripts and endogenous transcripts (**Figure S9A**). We specifically leveraged the presence of the synonymous single-nucleotide variant NM_004316.4(ASCL1);c.627C>G encoded in the ASCL1 lentivirus to distinguish exogenous and endogenous reads. The introduction of either exogenous ASCL1 or NeuroD1 to PRNB or PRNBS resulted in the induction of endogenous *ASCL1* expression (**Figure S9B**). This was also confirmed by the presence of reads mapping to the 5’ and 3’ untranslated regions (UTRs) which are present in endogenous transcripts but not encoded in exogenous proviral transcripts. Using a similar strategy of mapping reads to the 5’ and 3’ UTR, we also found that exogeneous NeuroD1 could activate endogenous *NEUROD1* expression (**Figure S9C**).

We then investigated how the different conditions in our NE reprogramming assay may impact the establishment of a transcriptional regulatory network (TRN) underlying an NEPC lineage state. We first focused on the differential expression of transcription factors due to SRRM4 (PRNBS vs. PRNB), ASCL1 (PRNBSA vs. PRNBS), and NeuroD1 (PRNBSN vs. PRNBS). We observed a significant induction of NE-associated transcription factors including INSM1, SOX2, SOX4, NK2 homeobox 2 (NKX2-2), and FOXA2 related to ASCL1 and NeuroD1 (**Figure 7A**). In stark contrast, not a single transcription factor demonstrated statistically significant differential expression upon the addition of SRRM4 to PRNB. Consistent with these findings, footprinting analysis of ATAC-seq data identified a significant enrichment in binding motifs associated with these NE-associated transcription factors in conditions of enforced ASCL1 and NeuroD1 expression but not SRRM4 expression (**Figure 7B**). Global chromatin accessibility profiles are also informative as they can differentiate lineage states in normal tissues and cancer. When we projected pseudo-bulk scATAC-seq data of reprogrammed C4-2B lines on the PCA space defined by ATAC-seq data of previously published ARPC and NEPC PDX models ^22^, we found that C4-2B lines reprogrammed with PRNBSA and PRNBSN clustered with the NEPC PDXs while those reprogrammed with PRNB and PRNBS were more similar to ARPC PDXs (**Figure 7C**). Further, we found that the expression of NEPC-associated cell surface targets (*DLL3*, *L1CAM*, *CEACAM5*, *RET* and *SEZ6*) were significantly increased and the ARPC-associated cell surface targets (*FOLH1, PSCA, STEAP1*, and *TACSTD2*) were significantly decreased upon the addition of ASCL1 or NeuroD1. In contrast, SRRM4 alone did not have a prominent effect on the expression of these cell surface targets (**Figure 7D**). Taken together, these findings indicate that, unlike ASCL1 and NeuroD1, SRRM4 activity and the resultant loss of REST repressor activity alone may be insufficient to drive a NEPC lineage due to the absence of active transcriptional and epigenetic reprogramming.

**Figure 7.**
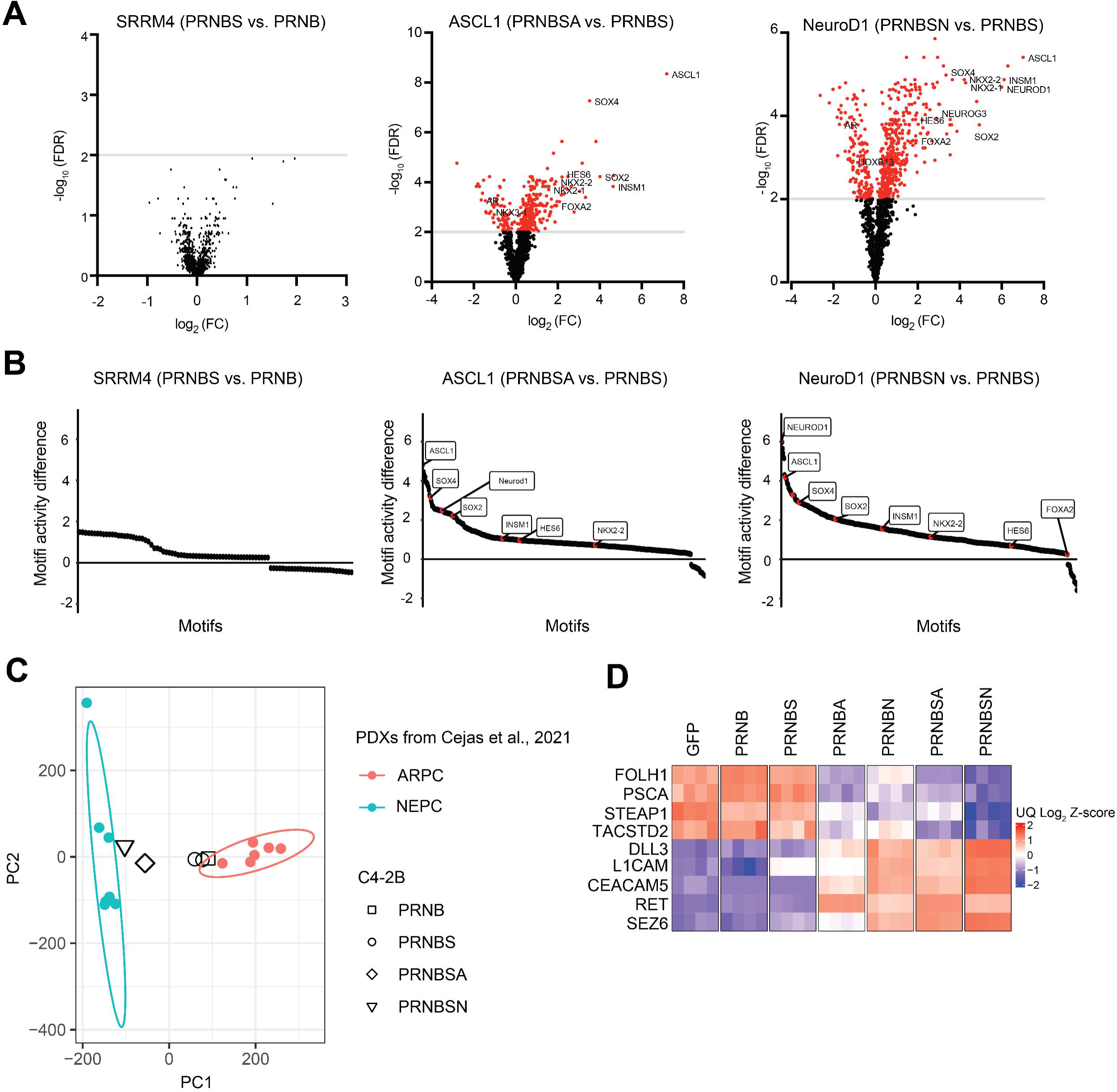
ASCL1/NeuroD1 induce the expression and activity of NE-associated transcription factors. (**A**) Volcano plots showing the differential expression of genes encoding transcriptions factors in pairwise comparisons of RNA-seq gene expression data of C4-2B cells reprogrammed with PRNBS vs. PRNB, PRNBSA vs. PRNBS, and PRNBSN vs. PRNBS. Red dots represent genes with a −log_10_(FDR) > 2. FC represents fold-change. (**B**) Plots of differential transcription factor motif activity from footprinting analysis of pseudo-bulk scATAC-seq data from C4-2B cells reprogrammed with PRNBS vs. PRNB, PRNBSA vs. PRNBS, and PRNBSN vs. PRNBS. Note that labelled motifs in the second and third panel do not show up in the first panel. (**C**) PCA of chromatin accessibility signals from pseudo-bulk scATAC-seq of reprogrammed C4-2B cell line conditions projected on previously published ATAC-seq data from NEPC and ARPC PDX models. (**D**) Heatmap of RNA-seq gene expression data showing cell surface targets relevant to ARPC and NEPC in reprogrammed C4-2B cell conditions.

### Downregulation of major histocompatibility complex class I (MHC I) in NEPC is functionally attributable to ASCL1/NeuroD1-driven programs

The downregulation of MHC I antigen presentation machinery by cancer to overcome immune surveillance and avoid immune destruction is an established hallmark of cancer. In particular, this phenomenon has been associated with cancers with neuroendocrine differentiation ^55^. In a recent study, examination of SCLC data sets showed that tumors with low NE scores demonstrated higher expression of MHC I genes while those with high NE scores showed decreased MHC I and reduced immune infiltration ^56^. Further, cancer cells are capable of co-opting PRC2 in a lineage-specific fashion to silence the MHC I antigen processing and presentation pathway ^57^. Mechanistically, mutation of the catalytic domain or pharmacologic inhibition of the PRC2 core subunit EZH2 was shown to impair MHC I repression, highlighting the critical role of PRC2 in this process.

We therefore asked whether a similar phenomenon may be operative in NEPC and could be functionally attributable to lineage-defining ASCL1 and/or NeuroD1 expression. We first examined a published RNA-seq gene expression dataset of mCRPC ^42^ including CRPC-Ad (adenocarcinoma) and CRPC-NE (NEPC) patient specimens and found that NEPC was associated with reduced MHC I antigen processing and presentation gene signature scores as well as lower *B2M* (beta-2-microglobulin, a critical component of MHC I) expression (**Figures 8A and S10**). A similar analysis using published data from tumors in the UW TAN PC rapid autopsy cohort also showed a reduced overall expression of *B2M* in the NEPC samples (**Figure 8B**). We then evaluated RNA-seq gene expression data from our reprogrammed C4-2B lines and correlated each condition and NE signature score with MHC I antigen processing and presentation gene signature scores. Consistent with prior literature, we discovered a reduction in MHC I gene expression with increasing NE differentiation (**Figure 8C**). Downregulation of MHC I gene expression started as early as D2 along with AR pathway repression (**Figure S11**). Furthermore, the addition of SRRM4 to PRNB did not have an appreciable effect on MHC I expression, indicating that loss of REST does not itself contribute to the downregulation of MHC I. In contrast, conditions where ASCL1 or NeuroD1 were added did demonstrate reduced expression of MHC I gene sets. We found that *B2M* expression was especially downregulated in conditions reprogrammed with NeuroD1 which were also associated with prominent NE signature scores (**Figure 8C**). These findings were in line with integrated scRNA-seq and scATAC-seq data from reprogrammed C4-2B cells showing an inverse correlation between NEPC score and *B2M* expression (**Figure 8D-F**) To confirm these findings at the protein level, we performed flow cytometry on reprogrammed C4-2B cells and showed that the PRNBSN condition was associated with a significant reduction in B2M positivity relative to the GFP control condition (29.2% vs. 60.9%) (**Figure 8G**). In addition, we observed that cells with diminished MHC I gene expression including B2M showed high EZH2 expression in both C4-2B and MDA PCa 2b cell lines (**Figure 8C and Figure S12**). Together, these findings functionally attribute the lineage-specific downregulation of MHC I in NEPC (relative to ARPC) to the lineage-defining transcription factors ASCL1 and NeuroD1.

**Figure 8.**
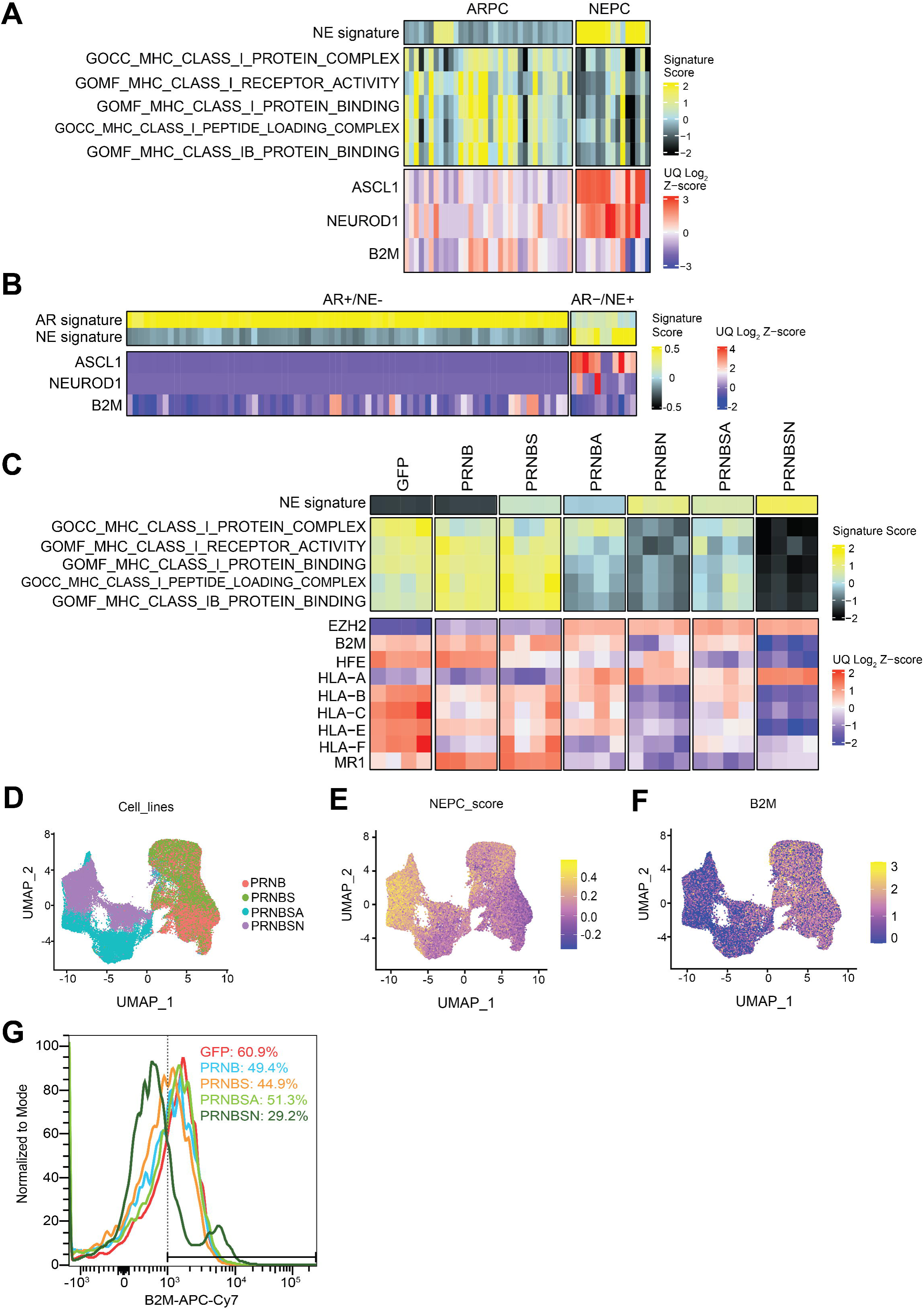
Downregulation of MHC class I antigen processing and presentation genes in NEPC is functionally attributable to ASCL1/NeuroD1. (**A**) Heatmap of RNA-seq gene expression data showing the NE signature and MHC class I pathway scores (top) and select genes (bottom) in ARPC and NEPC from Beltran et al., 2016. (**B**) Heatmap of RNA-seq gene expression data showing the AR and NE signature scores and select genes in AR^+^/NE^-^ and AR^-^/NE^+^ metastatic prostate cancers from the University of Washington Tissue Acquisition Necropsy (UW TAN) program. (**C**) Heatmap of RNA-seq gene expression data from reprogrammed C4-2B cell line conditions showing the NE signature and MHC class I pathway scores (top) and select MHC class I genes including B2M (bottom). UQ: upper quartile normalization. (**D**) UMAP analysis of scRNA-seq and scATAC-seq data from C4-2B cells reprogrammed with PRNB, PRNBS, PRNBSA or PRNBSN using the Seurat package. UMAP plots of reprogrammed cells colored based on (**E**) NEPC score and (**F**) B2M gene expression. (**G**) Flow cytometry histogram plots showing the percentage of B2M positive cells in reprogrammed C4-2B cells.

## DISCUSSION

AR signaling is pivotal in defining the luminal epithelial lineage of most PCs. However, potent therapies have been developed that apply stringent selective pressure against AR signaling, leading to the increased therapy-driven clinical presentation of lineage plasticity in PC. One such example is the conversion of ARPC to NEPC where the epithelial cancer state enforced by the AR program is replaced by a neural/NE cancer lineage ^7^. Alternative AR-null PC lineages have also been identified including double-negative prostate cancer (DNPC) which lacks both AR and NE marker expression ^18,58^. Further, this lineage plasticity in the setting of targeted therapies has also been observed in other epithelial cancer types such as epidermal growth factor receptor (EGFR)-mutant lung adenocarcinoma in which transformation to SCLC occurs in up to 15% of patients treated with EGFR tyrosine kinase inhibitors ^59^. However, the mechanisms by which AR (or EGFR) expression and signaling are effectively silenced during this transition have been poorly understood. The ability to functionally characterize these dynamic processes in a defined manner by reverse genetics has been limited by the lack of available model systems allowing for rapid manipulation and repeated sampling over time. Furthermore, due to documented heterogeneity, the application of narrow phenotypic markers to define NEPC has also been problematic and complicated our understanding of this PC lineage state.

We established an *in vitro* system in which we can reproducibly and durably reprogram established human ARPC cell lines to NEPC with defined genetic factors over a specified time course. This tractable system recapitulates key features of the process of NEtD including global, dynamic transcriptional and epigenetic reprogramming. Importantly, we uncovered the ability of the pioneer neural transcription factors ASCL1 and NeuroD1 to suppress AR expression and signaling by chromatin remodeling of the somatically acquired enhancer of AR and global ARBS with enhancer activity. These findings show that the activity of a new lineage-defining transcription factor can concurrently silence the prior cancer lineage program while establishing a divergent lineage. An immediate question that emerges from these findings is whether AR and ASCL1 or NeuroD1 activity is mutually exclusive in individual PC cells. Emerging single-cell sequencing data sets of human and mouse models of NEPC indicate that this may be the case ^60,61^. The competence of ASCL1 and/or NeuroD1 to silence the AR program in prostate cancer may be a major constraint to NEtD, providing a potential basis for why NEPC only occurs in 15% of therapy-driven mCRPC cases. How the expression of these neural/NE transcription factors is initially induced in PC also remains to be determined and warrants further investigation.

Our studies have also distinguished the relative contributions of SRRM4 and ASCL1 or NeuroD1 to NE reprogramming of PC, and yield insight into the problematic nature of using sparse protein (IHC) or RNA analysis based on only a few markers to define NEPC lineage states. In our reprogramming studies, we find that SRRM4 activity results in the loss of REST which activates neural/NE gene expression including SYP and CHGA. However, unlike ASCL1 or NeuroD1, SRRM4 does not induce a new transcriptional regulatory network based on transcription factor expression and inferred transcription factor activity analyses, does not elicit epigenomic shifts consistent with NEtD, does not affect AR expression and signaling, and does not alter the expression of ARPC-associated cell surface targets. Thus, our findings indicate that a reliance on a sparse set of NE markers, including highly canonical markers, to define NEPC can be flawed as their expression may be independent of true NEPC lineage states. A major clinical consideration is that mCRPCs that express select NE markers but have not undergone NEtD may still be responsive to therapies intended for ARPC including ARSIs and PSMA radioligand therapy.

In our NE reprogramming assay, we have found that ASCL1 and NeuroD1 but not other NE-associated transcription factors are competent to induce NEtD of PC. This supports a transcription regulatory network model where ASCL1 and NeuroD1 sit atop a hierarchy of NE-associated transcription factors necessary to establish self-sustaining NEPC transcriptional convergence. We have shown that ASCL1 and NeuroD1 are capable of self-regulation but the full set of regulatory interactions between these and other NE-associated transcription factors are unknown in NEPC. Future studies to define these relationships through genetic loss-of-function screening using this and other NEtD assays elucidate these functional interactions, identify potential vulnerabilities, and promote therapeutic development to inhibit NEtD under the pressures of otherwise effective targeted therapies.

Prior work identified suppression of MHC I antigen processing and presentation in neuroendocrine cancers and a functional association with lineage-specific PRC2 activity ^55–57^. Because our studies directly implicated ASCL1 and NeuroD1 in lineage reprogramming of ARPC to NEPC as well as the induction of PRC2 activity, we examined the expression of MHC I pathway gene sets and discovered that MHC I pathway genes were suppressed in the effective ASCL1- or NeuroD1-driven reprogramming conditions. In contrast, the addition of SRRM4 and subsequent loss of REST activity had little to no effect on the MHC I pathway. These findings support that downregulation of the MHC I antigen processing and presentation pathway in NEPC is functionally attributable to ASCL1 and NeuroD1 activity. Immune checkpoint inhibitors have demonstrated limited utility in mCRPC except in the case of DNA mismatch repair deficiency ^62^. Thus, our results support therapeutic efforts to induce immunogenic responses in NEPC using combinations with epigenetic inhibitors such as those targeting EZH2, histone deacetylases, or demethylases to enhance MHC I expression ^63,64^.

Overall, this study provides important mechanistic details of NEtD of PC by highlighting key lineage-defining activities of neural pioneer transcription factors in this process. These findings have important implications on the molecular definition of NEPC and therapeutic considerations related to the application of targeted agents and immune-based therapies.

## METHODS

### Cell lines

LNCaP (Cat# CRL-1740, RRID: CVCL_0395), C4-2B (Cat# CRL-3315, RRID: CVCL_4784), MDA PCa 2b (Cat# CRL-2422, RRID: CVCL_4748) cell lines were purchased from ATCC. All cell lines were validated by short tandem repeat analysis after receipt and on a yearly basis. LNCaP and C4-2B were maintained in RPMI medium supplemented with 10% FBS, 100 U/mL penicillin and 100 μg/mL streptomycin, and 4 mmol/L GlutaMAX. MDA PCa 2b was maintained in F-12K medium supplemented with 20% FBS, 25 ng/ml cholera toxin, 10 ng/ml mouse epidermal growth factor, 0.005 mM phosphoethanolamine, 100 pg/ml hydrocortisone, 1X insulin-transferrin-selenium (ITS-G), 100 U/mL penicillin and 100 μg/mL streptomycin, and 4 mmol/L GlutaMAX.

### Lentiviral constructs and lentivirus production

Double-barcoded lentiviral vectors used as a lentiviral backbone vector were generated from FU-CGW by sequentially inserting matched 10-nucleotide barcodes into the Pacl site distal to the HIV FLAP using the Quick Ligation Kit (New England Biolabs) and PCR amplification of the WPRE sequence and barcode with insertion into the Clal site proximal to the 3’ LTR by HiFi DNA Assembly (New England Biolabs). U6 promoter and shRNA cassettes were isolated by digesting pLKO.1 TRC shRNA clones with PspXI and EcorRI and were inserted into the digested double-barcoded plasmid as previously described ^8,9^. Lentiviral vectors expressing dominant-negative TP53, BCL2, MYC, SRRM4, NR0B2, KRAS, ASCL1 and NeuroD1 were generated by PCR amplification of the open reading frames and subcloning into the EcoRI site of double-barcoded lentiviral vectors by NEBuilder HiFi DNA Assembly (New England Biolabs). Lentiviruses were prepared and titered as previously described ^8^.

### Immunoblotting

Whole cell extracts were collected in 9 M urea lysis buffer and quantified using the Pierce Rapid Gold BCA Protein Assay Kit (Thermo Fisher Scientific). Protein samples were fractionated by SDS-PAGE using Bolt Bis-Tris Plus gels and transferred to a nitrocellulose membrane using Bolt Transfer Buffer according to the manufacturer’s instructions (Invitrogen). Membranes were blocked with 5% non-fat milk in PBST (DPBS + 0.5% Tween 20) for 30 minutes while shaking at room temperature, then incubated with primary antibodies at 4°C overnight. Membranes were washed three times for 5 minutes with PBST and incubated with horseradish peroxidase (HRP) conjugated anti-mouse or anti-rabbit secondary antibodies for 1 hour at room temperature. Blots were washed three times for 5 minutes each with PBST and developed with Immobilon Western Chemiluminescent HRP Substrate (EMD Millipore) for 3 minutes at room temperature. Blot images were acquired using a ChemiDoc Imaging System (Bio-Rad).

### Immunohistochemistry (IHC) and immunofluorescence (IF) studies

Cells and xenografts were fixed in 10% buffered formalin for 12 hours and then embedded in HistoGel (Thermo Fisher Scientific) and paraffin, sectioned, and mounted on glass slides (Thermo Fisher Scientific). IHC was performed as previously described ^26^. In brief, slides containing formalin-fixed paraffin-embedded sections were deparaffinized in xylene and rehydrated in 100%, 95%, and 70% ethanol. Slides were then heated in antigen retrieval buffer (0.2 mol/L citric acid and 0.2 mol/L sodium citrate) within a pressure cooker followed by PBS wash. Slides were blocked with 2.5% horse serum for 30 minutes and then incubated with primary antibody diluted in 2.5% horse serum overnight at 4°C. HRP was detected with ImmPRESS-HRP anti-mouse or anti-rabbit IgG peroxidase detection kits (Vector Laboratories) and staining was visualized with DAB peroxidase substrate (Dako). Slides were counterstained with hematoxylin and dehydrated for mounting.

Immunofluorescence studies using specific antibodies were carried out on archival formalin fixed paraffin embedded tissues. In brief, 5 µm paraffin sections were de-waxed and rehydrated following standard protocols. Antigen retrieval consisted of steaming for 40 minutes in Target Retrieval Solution (Agilent). Slides were then washed and equilibrated in TBS-Tween buffer (Sigma) for 10 minutes. Primary antibodies were applied at at 37°C for 60 minutes. Sequential dual-immunofluorescence labeling was carried out using Tyramide SuperBoost kits (Thermo Fisher).

### Neuroendocrine transdifferentiation assay

Cells were seeded in 6-well tissue culture plates at a density of 3 x 10^5^ cells per mL in 3 mL of RPMI medium supplemented with 10% FBS, 100 U/mL penicillin and 100 μg/mL streptomycin, and 4 mmol/L GlutaMAX. Cells were transduced approximately 4-6 hours after seeding at a defined multiplicity of infection (MOI) of 4 for each lentivirus. 72 hours after transduction, cells were trypsinized, washed, and transferred to 100 mm tissue culture plates in 15 mL of neural stem cell media (N-SCM) consisting of Advanced DMEM/F12 medium supplemented with 1X serum-free B27, 10 ng/mL recombinant human bFGF, 10 ng/mL recombinant human EGF, 100 U/mL penicillin and 100 μg/mL streptomycin, and 4 mmol/L GlutaMAX. Media were replenished every 3-4 days. Cells were collected 14 days post-transduction for analysis.

### Mouse studies

All animal care and studies were performed in accordance with an approved Fred Hutchinson Cancer Center Institutional Animal Care and Use Committee protocol and Comparative Medicine regulations. Eight-week-old male NSG (NOD-SCID-IL2Rγ-null, RRID:BCBC_4142) mice were obtained from the Jackson Laboratory. A total of 1 x 10^6^ cells from each C4-2B reprogramming condition were suspended in 100 μL of cold Matrigel (Corning) and implanted by injection subcutaneously into NSG mice.

### Single-cell DNA amplicon sequencing library preparation and sequencing

A custom panel was designed for the Mission Bio Tapestri to amplify segments of ten mouse genes at two exons each, the 5’ and 3’ lentiviral barcodes, and lentiviral GFP. Libraries were generated from C4-2B cells transduced with PRNBSA LV pool at MOI of 4 for each lentivirus using the Mission Bio Tapestri Single-cell DNA Custom Kit according to the manufacturer’s recommendations. Single cells (3,000 to 4,000 cells per μl) were resuspended in Tapestri cell buffer and encapsulated using a Tapestri microfluidics cartridge, lysed, and barcoded. Barcoded samples were subjected to targted PCR amplification and PCR products were removed from individual droplets, purified with KAPA Pure Beads (Roche Molecular Systems), and used as a template for PCR to incorporate Illumina P7 indices. PCR products were purified by KAPA Pure Beads, and quantified by Qubit dsDNA High Sensitivity Assay (ThermoFisher Scientific). Sample quality was assessed by Agilent TapeStation analysis. Libraries were pooled and sequenced on an Illumina MiSeq or HiSeq 2500 with 150 bp paired-end reads in the Fred Hutchinson Cancer Center Genomics Shared Resource.

### Cell proliferation assay

For cell proliferation assays, the CellTiter-Glo® 2.0 Cell Viability Assay system (Promega) was used according to the manufacturer’s instructions. 5□×□10^4^ cells were plated into each well in a 96-well and 25□μL per well of CellTiter-Glo® 2.0 reagent was added at each endpoint for analysis. After a 30 minute incubation at room temperature, luminescence was measured using a BioTek Synergy H1 Multimode Reader (Agilent). Six replicate wells per time point were used to obtain measures of cell proliferation.

### RNA-seq

Total RNA was extracted with a Purelink^TM^ RNA mini kit (Invitrogen). cDNA libraries were prepared from isolated RNA using the Illumina Stranded mRNA prep kit. High-throughput sequencing with 50 bp paired-end reads was performed using an Illumina NovaSeq 6000 SP. Sequencing reads were mapped to human genome reference hg38 and gene expression was quantified and normalized using the Toil RNA-seq pipeline (v4.1.2). Sequence adapters were trimmed using CutAdapt (v1.9) ^65^. Trimmed sequences were then aligned to human reference genome GRCh38 using STAR (v2.4.2a) ^66^ and gene expression quantification was performed using RSEM v1.2.25 ^67^. RSEM expected counts were subsequently normalized by upper-quartile (UQ) normalization for downstream analysis.

The enrichment scores were calculated in R using the GSVA package using a previously published set of 8 AR-regulated genes for AR-activity scores and the 14 genes in the NEURO I and NEURO II gene sets for NEURO scores^18^. Differential gene expression analysis was performed using the R package DEseq2 (v1.36.0) RNA-seq data from Beltran et al., 2016 was processed through the Toil RNA-seq pipeline (v4.1.2). Gene expression data was projected onto the Beltran et al., 2016 PCA space to examine reprogramming towards the NEPC phenotype. Gene signature scores were calculated based on RNA-seq expression data using the R package GSVA ^68^. The neuroendocrine score was previously described based on CRPC patient data ^42^. Briefly, 70 genes were used to construct the score based on differential expression or methylation between CRPC-NE and CRPC-Ad or relevance to CRPC-NE based on the literature. The mean expression of the 70 genes was calculated to form the NE score. The scores were visualized in heatmap format using ComplexHeatmap (v2.12.0) ^69^.

### Differential RNA alternative splicing analysis

Differential RNA alternative splicing (AS) between vector transduction conditions was calculated from bulk RNA-sequencing using the rMATS (version 4.1.1)^70^. Gene and transcript annotations were sourced from Ensembl GRCH38 release 104^71^. The program was run under the statoff mode to obtain the percent spliced in (PSI) matrix for all C42B samples. AS events that were defined by both junction counts and exon counts were used for downstream analysis. The PSI values of all five AS events: skipped exon (SE), alternative 5’ splice site (A5SS), alternative 3’ splice site (A3SS), mutually exclusive exons (MXE) and retained intron (RI) were aggregated together to form a large matrix for PCA. CRPC-Adeno and CRPC-NE Beltran et al 2016 patient RNAseq data was processed using the same rMATS workflow for AS calling. C42B was projected onto Betlran et al 2016 PSI PCA space for examining the remodeling of AS by oncogene transduction towards the NE direction.

### CUT&RUN

Cells were harvested by centrifugation at 1200 rpm for 3 minutes in a swinging bucket rotor and washed twice in ice cold wash buffer (20 mM HEPES pH 7.5; 150 mM NaCl; 0.5 mM Spermidine; 1 tablet of Roche complete EDTA-free protease inhibitor). Concanavalin A-coated beads (Bangs Laboratories) were activated by resuspension and washed in binding buffer (20 mM HEPES-KOH at pH 7.9; 10 mM KCl, 1 mM CaCl_2_; 1 mM MnCl_2_). Cells were mixed with activated beads at a ratio of 9:1 at room temperature for 10 minutes. The cell and bead mixtures were resuspended in antibody solution (2 mM EDTA; 0.05% digitonin in wash buffer). Corresponding antibodies were added at desired concentrations to 150 µl of cell and bead mixture and rotated overnight at 4°C. Libraries were prepared using the AutoCUT&RUN protocol (https://www.protocols.io/view/autocut-run-genome-wide-profiling-of-chromatin-pro-6qpvre6zblmk/v1). High-throughput sequencing with 50 bp paired-end reads was performed on CUT&RUN libraries using an Illumina NextSeq.

### Single-cell multiome RNA-seq and ATAC-seq

Cells were collected 14 days post transduction and processed per manufacturer’s instructions using the 10x Chromium Next GEM Single Cell Multiome ATAC + Gene Expression platform to generate libraries. Single-cell RNA-seq libraries and single-cell ATAC-seq libraries were sequenced on an Illumina NovaSeq 6000 targeting 35,000 read pairs/nuclei for RNA-seq and 40,000 read pairs/nucleus for ATAC-seq.

### Single-cell RNA-seq and ATAC-seq analysis

Single-cell multiome ATAC- and RNA-sequencing reads were processed through the CellRanger ARC analysis pipeline. RNA-sequencing filtered feature count matrices and ATAC-sequencing fragment paths from each sample were loaded individually loaded into R with Signac (version 1.7.0)^72^. The data was filtered for total RNA count between 1000 and 50000, total feature RNA count between 500 and 10000, percent mitochondria RNA less than 20%, total ATAC fragment count between 1000 and 50000, nucleosome signal score less than 2, and a TSS enrichment score above 1.5. ATACseq peaks were called using MACS2 ^73^. Peaks on nonstandard chromosomes and in genomic blacklist regions of hg38 were removed. ATAC counts were quantified for all peaks across all samples.

### Single-cell trajectory analysis

Filtered Seurat object was loaded into Monocle3 (version 1.2.9) to calculate Single-cell trajectory ^74,75^. First 50 PCA dimensions were used in the preprocessing step. Cells were aligned by regressing on percent mitochondria RNA.

### CUT&RUN data processing and analyses

Peak calls for H3K4me1, H3K4me3 and H3K27ac from CUT&RUN data at different timepoints were performed using SEACR (v1.3) ^76^. Bedgraph files were prepared using bedtools (v2.27.1) to serve as the inputs for SEACR. IgG control conditions at each timepoint were used to define empirical thresholds for peak calling. The “norm” parameter was used in SEACR to normalize the data. Peaks were called in “stringent” mode. Active promoters were defined by checking if H3K4me3 peaks overlap with H3K27ac signals using the bedtools intersect -wa option. The resulting regions within +/- 5 kb of transcription start sites (TSS) were retained. TSS regions were calculated by applying GTFtools (v0.8.5) ^77^ to the gencode v39 (GRCh38.p13) GTF file. Active enhancers were obtained by overlapping H3K4me1 peaks with H3K27ac peaks, followed by subtracting the +/- 5 kb TSS regions using the bedtools subtract –A option. Super-enhancers were called using ROSE (v0.1) ^52,78^ on H3K27ac CUT&RUN data. STITCHING_DISTANCE was set to 12.5 kb. TSS_EXCLUSION_ZONE_SIZE was set to 2.5 kb. hg38 was used as the reference genome. The resulting genomic regions of active promoters, active enhancers and super-enhancers were loaded into DiffBind ^79^ for differential binding analysis. Counts that mapped to the genomic regions were normalized using the Relative Log Expression (RLE) method. For super-enhancers, the counts of member regions that belong to each parental broad genomic region were aggregated together to form the overall count. The correlations between time and genomic regions were calculated using Kendall test followed by Benjamini-Hochberg P-value adjustment. Genomic regions were mapped to genes using ChIPseeker (v1.32.0) ^80^. Genomic regions and genes demonstrating significant changes along with time were visualized in heatmap format using ComplexHeatmaps (v2.12.0)^69^ and analyzed by Gene Ontology using hypergeometric test with MSigDB gene sets ^81^.

### AR binding site analyses

Inducible, constitutively active, and inactive AR binding sites (ARBS) were obtained from a previously published study ^53^. A random set of genomic regions with the same length of ARBS (700 bp) was generated by randomly selecting 50 start sites for each chromosome. Fragments of scATAC-seq data were quantified against these sites. The significance of the difference between conditions was calculated based on the log transformed counts using Welch’s t-test.

### AR enhancer analyses

The genomic coordinates of the somatically acquired AR enhancer were obtained from a previously published study ^5^. scATAC-seq data were examined at the same overall genomic region to determine the location of the AR enhancer in our model system and in the prostate cancer hepatic metastasis. The correlation between scATAC chromatin accessibility and scRNA gene expression in reprogrammed cells was visualized using CoveragePlot in the R package Signac (version 1.7.0) ^72^.

### Statistical analyses

Data analysis was performed on GraphPad Prism 9 (GraphPad Software, Inc.). Quantitative PCR results were analyzed in Excel. Statistical significance was determined using the unpaired two-tailed Student t test, unpaired two-tailed Welch t test where the variances are shown to be different via F-test, one-way ANOVA, or two-way ANOVA. Only two-tailed tests were used. Results are depicted as mean + SD unless stated otherwise. All P values of <0.05, <0.01, <0.001, and <0.0001 were considered significant. Pearson correlation coefficient was used to determine correlation between genes. The symbols used to represent the P values were: ns, nonsignificant for P > 0.05; *, P ≤ 0.05; **, P ≤ 0.01; ***, P ≤ 0.001; ****, P ≤ 0.0001. The test used in each statistical analysis is specified in the figure legends.

## Data availability

All raw and analyzed sequencing data including RNA-seq, scRNA-seq, and scATAC-seq have been uploaded to the National Center for Biotechnology Information (NCBI) Gene Expression Omnibus (GEO) under accession SuperSeries GSE225026 and are publicly available for download.

## ACKNOWLEDGEMENTS

We would like to thank the Fred Hutch Genomics Shared Resource, Comparative Medicine Shared Resource, and Experimental Histopathology Shared Resource (supported by NIH/NCI Cancer Center Support Grant P30CA015704). We also acknowledge support by the Pacific Northwest Prostate Cancer SPORE (NCI P50CA97186) and the Institute for Prostate Cancer Research. SL was supported by a Department of Defense Prostate Cancer Research Program Early Investigator Research Award (W81XWH-20-1-0083). The work was also supported by NIH DP2 CA271301 (J.K.L.); NCI R01 CA234715 (J.K.L., M.C.H., P.S.N.); NCI R01 CA266452 (J.K.L., P.S.N., M.C.H.); NCI R01 CA280056 (P.S.N.); NCI R01 CA222877 (T.G.G.); NCI P50CA092131 (T.G.G., J.K.L.); NCI P01CA163227 (P.S.N., H.W.L.); the W.M. Keck Foundation (T.G.G.); the V Foundation (M.C.H.); the UCLA Eli and Edythe Broad Center of Regenerative Medicine and Stem Cell Research (T.G.G.); Hal Gaba Director’s Fund for Cancer Stem Cell Research (T.G.G.); and Prostate Cancer Foundation Challenge Awards (J.K.L., P.S.N.).

## AUTHOR CONTRIBUTIONS

Conceptualization, S.L., K.S., H.S., T.G.G., and J.K.L.; Methodology, S.L. and J.K.L.; Investigation, S.L., K.S., H.S., Y.T., A.H., V.B., B.H., R.A.P., H.W.L., C.M., M.C.H.; Writing – Original Draft, S.L. and J.K.L.; Writing – Review & Editing, P.S.N., T.G.G., and J.K.L.; Resources, C.M., P.S.N.; Supervision, T.G.G. and J.K.L.

## REFERENCES

1. Chen, C.D., Welsbie, D.S., Tran, C., Baek, S.H., Chen, R., Vessella, R., Rosenfeld, M.G., and Sawyers, C.L. (2004). Molecular determinants of resistance to antiandrogen therapy. Nature Medicine 10, 33–39. 10.1038/nm972.

2. Watson, P.A., Arora, V.K., and Sawyers, C.L. (2015). Emerging mechanisms of resistance to androgen receptor inhibitors in prostate cancer. Nat Rev Cancer 15, 701–711. 10.1038/nrc4016.

3. Hu, R., Dunn, T.A., Wei, S.Z., Isharwal, S., Veltri, R.W., Humphreys, E., Han, M., Partin, A.W., Vessella, R.L., Isaacs, W.B., et al. (2009). Ligand-Independent Androgen Receptor Variants Derived from Splicing of Cryptic Exons Signify Hormone-Refractory Prostate Cancer. Cancer Research 69, 16–22. 10.1158/0008-5472.Can-08-2764.

4. Viswanathan, S.R., Ha, G., Hoff, A.M., Wala, J.A., Carrot-Zhang, J., Whelan, C.W., Haradhvala, N.J., Freeman, S.S., Reed, S.C., Rhoades, J., et al. (2018). Structural Alterations Driving Castration-Resistant Prostate Cancer Revealed by Linked-Read Genome Sequencing. Cell 174, 433-+. 10.1016/j.cell.2018.05.036.

5. Takeda, D.Y., Spisak, S., Seo, J.H., Bell, C., O’Connor, E., Korthauer, K., Ribli, D., Csabai, I., Solymosi, N., Szallasi, Z., et al. (2018). A Somatically Acquired Enhancer of the Androgen Receptor Is a Noncoding Driver in Advanced Prostate Cancer. Cell 174, 422–432 e413. 10.1016/j.cell.2018.05.037.

6. Beltran, H., Rickman, D.S., Park, K., Chae, S.S., Sboner, A., MacDonald, T.Y., Wang, Y., Sheikh, K.L., Terry, S., Tagawa, S.T., et al. (2011). Molecular characterization of neuroendocrine prostate cancer and identification of new drug targets. Cancer discovery 1, 487–495. 10.1158/2159-8290.Cd-11-0130.

7. Beltran, H., Hruszkewycz, A., Scher, H.I., Hildesheim, J., Isaacs, J., Yu, E.Y., Kelly, K., Lin, D., Dicker, A., Arnold, J., et al. (2019). The Role of Lineage Plasticity in Prostate Cancer Therapy Resistance. Clin Cancer Res 25, 6916–6924. 10.1158/1078-0432.Ccr-19-1423.

8. Lee, J.K., Phillips, J.W., Smith, B.A., Park, J.W., Stoyanova, T., McCaffrey, E.F., Baertsch, R., Sokolov, A., Meyerowitz, J.G., Mathis, C., et al. (2016). N-Myc Drives Neuroendocrine Prostate Cancer Initiated from Human Prostate Epithelial Cells. Cancer cell 29, 536–547. 10.1016/j.ccell.2016.03.001.

9. Park, J.W., Lee, J.K., Sheu, K.M., Wang, L., Balanis, N.G., Nguyen, K., Smith, B.A., Cheng, C., Tsai, B.L., Cheng, D., et al. (2018). Reprogramming normal human epithelial tissues to a common, lethal neuroendocrine cancer lineage. Science (New York, N.Y.) 362, 91–95. 10.1126/science.aat5749.

10. Nyquist, M.D., Corella, A., Coleman, I., De Sarkar, N., Kaipainen, A., Ha, G., Gulati, R., Ang, L., Chatterjee, P., Lucas, J., et al. (2020). Combined TP53 and RB1 Loss Promotes Prostate Cancer Resistance to a Spectrum of Therapeutics and Confers Vulnerability to Replication Stress. Cell Rep 31. ARTN 107669 10.1016/j.celrep.2020.107669.

11. Berger, A., Brady, N.J., Bareja, R., Robinson, B., Conteduca, V., Augello, M.A., Ruca, L., Ahmed, A., Dardenne, E., Lu, X.D., et al. (2019). N-Myc mediated epigenetic reprogramming drives lineage plasticity in advanced prostate cancer. J Clin Invest 129, 3924–3940. 10.1172/Jci127961.

12. Zhang, X.T., Coleman, I.M., Brown, L.G., True, L.D., Kollath, L., Lucas, J.M., Lam, H.M., Dumpit, R., Corey, E., Chery, L., et al. (2015). SRRM4 Expression and the Loss of REST Activity May Promote the Emergence of the Neuroendocrine Phenotype in Castration-Resistant Prostate Cancer. Clin Cancer Res 21, 4698–4708. 10.1158/1078-0432.Ccr-15-0157.

13. Mu, P., Zhang, Z., Benelli, M., Karthaus, W.R., Hoover, E., Chen, C.C., Wongvipat, J., Ku, S.Y., Gao, D., Cao, Z., et al. (2017). SOX2 promotes lineage plasticity and antiandrogen resistance in TP53- and RB1-deficient prostate cancer. Science (New York, N.Y.) 355, 84–88. 10.1126/science.aah4307.

14. Lin, D., Wyatt, A.W., Xue, H., Wang, Y., Dong, X., Haegert, A., Wu, R., Brahmbhatt, S., Mo, F., Jong, L., et al. (2014). High fidelity patient-derived xenografts for accelerating prostate cancer discovery and drug development. Cancer Res 74, 1272–1283. 10.1158/0008-5472.CAN-13-2921-T.

15. Akamatsu, S., Wyatt, Alexander W., Lin, D., Lysakowski, S., Zhang, F., Kim, S., Tse, C., Wang, K., Mo, F., Haegert, A., et al. (2015). The Placental Gene PEG10 Promotes Progression of Neuroendocrine Prostate Cancer. Cell reports 12, 922–936. 10.1016/j.celrep.2015.07.012.

16. Epstein, J.I., Amin, M.B., Beltran, H., Lotan, T.L., Mosquera, J.M., Reuter, V.E., Robinson, B.D., Troncoso, P., and Rubin, M.A. (2014). Proposed Morphologic Classification of Prostate Cancer With Neuroendocrine Differentiation. Am J Surg Pathol 38, 756–767. 10.1097/Pas.0000000000000208.

17. Balanis, N.G., Sheu, K.M., Esedebe, F.N., Patel, S.J., Smith, B.A., Park, J.W., Alhani, S., Gomperts, B.N., Huang, J., Witte, O.N., and Graeber, T.G. (2019). Pan-cancer Convergence to a Small-Cell Neuroendocrine Phenotype that Shares Susceptibilities with Hematological Malignancies. Cancer Cell 36, 17-+. 10.1016/j.ccell.2019.06.005.

18. Labrecque, M.P., Coleman, I.M., Brown, L.G., True, L.D., Kollath, L., Lakely, B., Nguyen, H.M., Yang, Y.C., da Costa, R.M.G., Kaipainen, A., et al. (2019). Molecular profiling stratifies diverse phenotypes of treatment-refractory metastatic castration-resistant prostate cancer. J Clin Invest 129, 4492–4505. 10.1172/Jci128212.

19. Merkens, L., Sailer, V., Lessel, D., Janzen, E., Greimeier, S., Kirfel, J., Perner, S., Pantel, K., Werner, S., and von Amsberg, G. (2022). Aggressive variants of prostate cancer: underlying mechanisms of neuroendocrine transdifferentiation. J Exp Clin Canc Res 41. ARTN 46 10.1186/s13046-022-02255-y.

20. Raposo, A.A.S.F., Vasconcelos, F.F., Drechsel, D., Marie, C., Johnston, C., Dolle, D., Bithell, A., Gillotin, S., van den Berg, D.L.C., Ettwiller, L., et al. (2015). Ascl1 Coordinately Regulates Gene Expression and the Chromatin Landscape during Neurogenesis. Cell Rep 10, 1544–1556. 10.1016/j.celrep.2015.02.025.

21. Borromeo, M.D., Savage, T.K., Kollipara, R.K., He, M., Augustyn, A., Osborne, J.K., Girard, L., Minna, J.D., Gazdar, A.F., Cobb, M.H., and Johnson, J.E. (2016). ASCL1 and NEUROD1 Reveal Heterogeneity in Pulmonary Neuroendocrine Tumors and Regulate Distinct Genetic Programs. Cell Rep 16, 1259–1272. 10.1016/j.celrep.2016.06.081.

22. Cejas, P., Xie, Y.T., Font-Tello, A., Lim, K., Syamala, S., Qiu, X.T., Tewari, A.K., Shah, N., Nguyen, H.M., Patel, R.A., et al. (2021). Subtype heterogeneity and epigenetic convergence in neuroendocrine prostate cancer. Nat Commun 12. ARTN 5775 10.1038/s41467-021-26042-z.

23. Costanzo, F., Diez, M.M., Nunez, G.S., Diaz-Hernandez, J.I., Robles, C.M.G., Perez, J.D., Compe, E., Ricci, R., Li, T.K., Coin, F., et al. (2022). Promoters of ASCL1-and NEUROD1-dependent genes are specific targets of lurbinectedin in SCLC cells. Embo Mol Med 14. ARTN e14841 10.15252/emmm.202114841.

24. Baine, M.K., Hsieh, M.S., Lai, W.V., Egger, J.V., Jungbluth, A.A., Daneshbod, Y., Beras, A., Spencer, R., Lopardo, J., Bodd, F., et al. (2020). SCLC Subtypes Defined by ASCL1, NEUROD1, POU2F3, and YAP1: A Comprehensive Immunohistochemical and Histopathologic Characterization. J Thorac Oncol 15, 1823–1835. 10.1016/j.jtho.2020.09.009.

25. Szeitz, B., Megyesfalvi, Z., Woldmar, N., Valko, Z., Schwendenwein, A., Barany, N., Paku, S., Laszlo, V., Kiss, H., Bugyik, E., et al. (2022). In-depth proteomic analysis reveals unique subtype-specific signatures in human small-cell lung cancer. Clin Transl Med 12. ARTN e1060 10.1002/ctm2.1060.

26. DeLucia, D.C., Cardillo, T.M., Ang, L., Labrecque, M.P., Zhang, A., Hopkins, J.E., De Sarkar, N., Coleman, I., da Costa, R.M.G., Corey, E., et al. (2021). Regulation of CEACAM5 and Therapeutic Efficacy of an Anti-CEACAM5-SN38 Antibody-drug Conjugate in Neuroendocrine Prostate Cancer. Clin Cancer Res 27, 759–774. 10.1158/1078-0432.CCR-20-3396.

27. Augustyn, A., Borromeo, M., Wang, T., Fujimoto, J., Shao, C., Dospoy, P.D., Lee, V., Tan, C., Sullivan, J.P., Larsen, J.E., et al. (2014). ASCL1 is a lineage oncogene providing therapeutic targets for high-grade neuroendocrine lung cancers. Proceedings of the National Academy of Sciences of the United States of America 111, 14788–14793. 10.1073/pnas.1410419111.

28. Nouruzi, S., Ganguli, D., Tabrizian, N., Kobelev, M., Sivak, O., Namekawa, T., Thaper, D., Baca, S.C., Freedman, M.L., Aguda, A., et al. (2022). ASCL1 activates neuronal stem cell-like lineage programming through remodeling of the chromatin landscape in prostate cancer. Nat Commun 13. ARTN 2282 10.1038/s41467-022-29963-5.

29. Ku, S.Y., Rosario, S., Wang, Y., Mu, P., Seshadri, M., Goodrich, Z.W., Goodrich, M.M., Labbe, D.P., Gomez, E.C., Wang, J., et al. (2017). Rb1 and Trp53 cooperate to suppress prostate cancer lineage plasticity, metastasis, and antiandrogen resistance. Science (New York, N.Y.) 355, 78–83. 10.1126/science.aah4199.

30. Dardenne, E., Beltran, H., Benelli, M., Gayvert, K., Berger, A., Puca, L., Cyrta, J., Sboner, A., Noorzad, Z., MacDonald, T., et al. (2016). N-Myc Induces an EZH2-Mediated Transcriptional Program Driving Neuroendocrine Prostate Cancer. Cancer cell 30, 563–577. 10.1016/j.ccell.2016.09.005.

31. Li, Y.A., Donmez, N., Sahinalp, C., Xie, N., Wang, Y.W., Xue, H., Mo, F., Beltran, H., Gleave, M., Wang, Y.Z., et al. (2017). SRRM4 Drives Neuroendocrine Transdifferentiation of Prostate Adenocarcinoma Under Androgen Receptor Pathway Inhibition. Eur Urol 71, 68–78. 10.1016/j.eururo.2016.04.028.

32. Gobinet, J., Auzou, G., Nicolas, J.C., Sultan, C., and Jalaguier, S. (2001). Characterization of the interaction between androgen receptor and a new transcriptional inhibitor, SHP. Biochemistry 40, 15369–15377. 10.1021/bi011384o.

33. Babos, K.N., Galloway, K.E., Kisler, K., Zitting, M., Li, Y.C., Shi, Y.X., Quintino, B., Chow, R.H., Zlokovic, B.V., and Ichida, J.K. (2019). Mitigating Antagonism between Transcription and Proliferation Allows Near-Deterministic Cellular Reprogramming. Cell Stem Cell 25, 486-+. 10.1016/j.stem.2019.08.005.

34. Jouravel, N., Sablin, E., Arnold, L.A., Guy, R.K., and Fletterick, R.J. (2007). Interaction between the androgen receptor and a segment of its corepressor SHP. Acta Crystallogr D 63, 1198–1200. 10.1107/S0907444907045702.

35. Judware, R., and Culp, L.A. (1995). Over-expression of transfected N-myc oncogene in human SKNSH neuroblastoma cells down-regulates expression of beta 1 integrin subunit. Oncogene 11, 2599–2607.

36. Cobrinik, D., Francis, R.O., Abramson, D.H., and Lee, T.C. (2006). Rb induces a proliferative arrest and curtails Brn-2 expression in retinoblastoma cells. Mol Cancer 5, 72. 10.1186/1476-4598-5-72.

37. Li, Y.N., Chen, R.Q., Bowden, M., Mo, F., Lin, Y.Y., Gleave, M., Collins, C., and Dong, X.S. (2017). Establishment of a neuroendocrine prostate cancer model driven by the RNA splicing factor SRRM4. Oncotarget 8, 66878–66888. 10.18632/oncotarget.19916.

38. Li, S., Wong, A., Sun, H., Bhatia, V., Javier, G., Jana, S., Montgomery, R.B., Wright, J.L., Lam, H.M., Hsieh, A.C., et al. (2023). Combinatorial genetic strategy accelerates the discovery of cancer genotype-phenotype associations. bioRxiv. 10.1101/2023.04.12.536652.

39. Vue, T.Y., Kollipara, R.K., Borromeo, M.D., Smith, T., Mashimo, T., Burns, D.K., Bachoo, R.M., and Johnson, J.E. (2020). ASCL1 regulates neurodevelopmental transcription factors and cell cycle genes in brain tumors of glioma mouse models. Glia 68, 2613–2630. 10.1002/glia.23873.

40. Han, M., Li, F., Zhang, Y., Dai, P., He, J., Li, Y., Zhu, Y., Zheng, J., Huang, H., Bai, F., and Gao, D. (2022). FOXA2 drives lineage plasticity and KIT pathway activation in neuroendocrine prostate cancer. Cancer cell 40, 1306–1323.e1308. 10.1016/j.ccell.2022.10.011.

41. Bishop, J.L., Thaper, D., Vahid, S., Davies, A., Ketola, K., Kuruma, H., Jama, R., Nip, K.M., Angeles, A., Johnson, F., et al. (2017). The Master Neural Transcription Factor BRN2 Is an Androgen Receptor-Suppressed Driver of Neuroendocrine Differentiation in Prostate Cancer. Cancer discovery 7, 54–71. 10.1158/2159-8290.Cd-15-1263.

42. Beltran, H., Prandi, D., Mosquera, J.M., Benelli, M., Puca, L., Cyrta, J., Marotz, C., Giannopoulou, E., Chakravarthi, B.V., Varambally, S., et al. (2016). Divergent clonal evolution of castration-resistant neuroendocrine prostate cancer. Nat Med 22, 298–305. 10.1038/nm.4045.

43. Ohnishi, T., Shirane, M., and Nakayama, K.I. (2017). SRRM4-dependent neuron-specific alternative splicing of protrudin transcripts regulates neurite outgrowth. Sci Rep-Uk 7. ARTN 41130 10.1038/srep41130.

44. Tsoi, J., Robert, L., Paraiso, K., Galvan, C., Sheu, K.M., Lay, J., Wong, D.J.L., Atefi, M., Shirazi, R., Wang, X.Y., et al. (2018). Multi-stage Differentiation Defines Melanoma Subtypes with Differential Vulnerability to Drug-Induced Iron-Dependent Oxidative Stress. Cancer Cell 33, 890-+. 10.1016/j.ccell.2018.03.017.

45. Richard, A., Boullu, L., Herbach, U., Bonnafoux, A., Morin, V., Vallin, E., Guillemin, A., Gao, N.P., Gunawan, R., Cosette, J., et al. (2016). Single-Cell-Based Analysis Highlights a Surge in Cell-to-Cell Molecular Variability Preceding Irreversible Commitment in a Differentiation Process. Plos Biol 14. ARTN e1002585 10.1371/journal.pbio.1002585.

46. Basili, D., Zhang, J.L., Herbert, J., Krol, K., Denslow, N.D., Martyniuk, C.J., Falciani, F., and Antczak, P. (2018). In Silico Computational Transcriptomics Reveals Novel Endocrine Disruptors in Largemouth Bass (Micropterus salmoides). Environ Sci Technol 52, 7553–7565. 10.1021/acs.est.8b02805.

47. Jiang, S., Williams, K., Kong, X.D., Zeng, W.H., Nguyen, N.V., Ma, X.Y., Tawil, R., Yokomori, K., and Mortazavi, A. (2020). Single-nucleus RNA-seq identifies divergent populations of FSHD2 myotube nuclei. Plos Genet 16. ARTN e1008754 10.1371/journal.pgen.1008754.

48. Karanikolas, B.D.W., Figueiredo, M.L., and Wu, L. (2010). Comprehensive Evaluation of the Role of EZH2 in the Growth, Invasion, and Aggression of a Panel of Prostate Cancer Cell Lines. Prostate 70, 675–688. 10.1002/pros.21112.

49. Cyrta, J., Augspach, A., De Filippo, M.R., Prandi, D., Thienger, P., Benelli, M., Cooley, V., Bareja, R., Wilkes, D., Chae, S.S., et al. (2020). Role of specialized composition of SWI/SNF complexes in prostate cancer lineage plasticity. Nat Commun 11. ARTN 5549 10.1038/s41467-020-19328-1.

50. Crispatzu, G., Rehimi, R., Pachano, T., Bleckwehl, T., Cruz-Molina, S., Xiao, C., Mahabir, E., Bazzi, H., and Rada-Iglesias, A. (2021). The chromatin, topological and regulatory properties of pluripotency-associated poised enhancers are conserved in vivo. Nat Commun 12. ARTN 4344 10.1038/s41467-021-24641-4.

51. Stepniak, K., Machnicka, M.A., Mieczkowski, J., Macioszek, A., Wojtas, B., Gielniewski, B., Poleszak, K., Perycz, M., Krol, S.K., Guzik, R., et al. (2021). Mapping chromatin accessibility and active regulatory elements reveals pathological mechanisms in human gliomas (vol 12, 3621, 2021). Nat Commun 12. ARTN 7218 10.1038/s41467-021-27448-5.

52. Whyte, W.A., Orlando, D.A., Hnisz, D., Abraham, B.J., Lin, C.Y., Kagey, M.H., Rahl, P.B., Lee, T.I., and Young, R.A. (2013). Master Transcription Factors and Mediator Establish Super-Enhancers at Key Cell Identity Genes. Cell 153, 307–319. 10.1016/j.cell.2013.03.035.

53. Huang, C.C.F., Lingadahalli, S., Morova, T., Ozturan, D., Hu, E., Yu, I.P.L., Linder, S., Hoogstraat, M., Stelloo, S., Sar, F., et al. (2021). Functional mapping of androgen receptor enhancer activity. Genome Biol 22. ARTN 149 10.1186/s13059-021-02339-6.

54. Saint-Andre, V., Federation, A.J., Lin, C.Y., Abraham, B.J., Reddy, J., Lee, T.I., Bradner, J.E., and Young, R.A. (2016). Models of human core transcriptional regulatory circuitries. Genome Res 26, 385–396. 10.1101/gr.197590.115.

55. Dhatchinamoorthy, K., Colbert, J.D., and Rock, K.L. (2021). Cancer Immune Evasion Through Loss of MHC Class I Antigen Presentation. Front Immunol 12. ARTN 636568 10.3389/fimmu.2021.636568.

56. Cai, L., Liu, H.Y., Huang, F., Fujimoto, J., Girard, L., Chen, J., Li, Y.W., Zhang, Y.A., Deb, D., Stastny, V., et al. (2021). Cell-autonomous immune gene expression is repressed in pulmonary neuroendocrine cells and small cell lung cancer. Commun Biol 4. ARTN 314 10.1038/s42003-021-01842-7.

57. Burr, M.L., Sparbier, C.E., Chan, K.L., Chan, Y.C., Kersbergen, A., Lam, E.Y.N., Azidis-Yates, E., Vassiliadis, D., Bell, C.C., Gilan, O., et al. (2019). An Evolutionarily Conserved Function of Polycomb Silences the MHC Class I Antigen Presentation Pathway and Enables Immune Evasion in Cancer. Cancer Cell 36, 385-+. 10.1016/j.ccell.2019.08.008.

58. Bluemn, E.G., Coleman, I.M., Lucas, J.M., Coleman, R.T., Hernandez-Lopez, S., Tharakan, R., Bianchi-Frias, D., Dumpit, R.F., Kaipainen, A., Corella, A.N., et al. (2017). Androgen Receptor Pathway-Independent Prostate Cancer Is Sustained through FGF Signaling. Cancer Cell 32, 474-+. 10.1016/j.ccell.2017.09.003.

59. Sequist, L.V., Waltman, B.A., Dias-Santagata, D., Digumarthy, S., Turke, A.B., Fidias, P., Bergethon, K., Shaw, A.T., Gettinger, S., Cosper, A.K., et al. (2011). Genotypic and Histological Evolution of Lung Cancers Acquiring Resistance to EGFR Inhibitors. Sci Transl Med 3. ARTN 75ra26 10.1126/scitranslmed.3002003.

60. Wang, Z.W., Wang, T., Hong, D.N., Dong, B.J., Wang, Y., Huang, H.Q., Zhang, W.H., Lian, B.J., Ji, B.Y., Shi, H.Q., et al. (2022). Single-cell transcriptional regulation and genetic evolution of neuroendocrine prostate cancer. Iscience 25. ARTN 104576 10.1016/j.isci.2022.104576.

61. Brady, N.J., Bagadion, A.M., Singh, R., Conteduca, V., Van Emmenis, L., Arceci, E., Pakula, H., Carelli, R., Khani, F., Bakht, M., et al. (2021). Temporal evolution of cellular heterogeneity during the progression to advanced AR-negative prostate cancer. Nature communications 12, 3372. 10.1038/s41467-021-23780-y.

62. Abida, W., Cheng, M.L., Armenia, J., Middha, S., Autio, K.A., Vargas, H.A., Rathkopf, D., Morris, M.J., Danila, D.C., Slovin, S.F., et al. (2019). Analysis of the Prevalence of Microsatellite Instability in Prostate Cancer and Response to Immune Checkpoint Blockade. JAMA oncology 5, 471–478. 10.1001/jamaoncol.2018.5801.

63. Hiatt, J.B., Sandborg, H., Garrison, S.M., Arnold, H.U., Liao, S.Y., Norton, J.P., Friesen, T.J., Wu, F.N., Sutherland, K.D., Rienhoff, H.Y., et al. (2022). Inhibition of LSD1 with Bomedemstat Sensitizes Small Cell Lung Cancer to Immune Checkpoint Blockade and T-Cell Killing. Clin Cancer Res 28, 4551–4564. 10.1158/1078-0432.Ccr-22-1128.

64. Nguyen, E.M., Taniguchi, H., Chan, J.M., Zhan, Y.Q.A., Chen, X.P., Qiu, J., de Stanchina, E., Allaj, V., Shah, N.S., Uddin, F., et al. (2022). Targeting Lysine-Specific Demethylase 1 Rescues Major Histocompatibility Complex Class I Antigen Presentation and Overcomes Programmed Death-Ligand 1 Blockade Resistance in SCLC. J Thorac Oncol 17, 1014–1031. 10.1016/j.jtho.2022.05.014.

65. Martin, M. (2011). Cutadapt removes adapter sequences from high-throughput sequencing reads. EMBnet.journal 17, 10–12.

66. Dobin, A., Davis, C.A., Schlesinger, F., Drenkow, J., Zaleski, C., Jha, S., Batut, P., Chaisson, M., and Gingeras, T.R. (2013). STAR: ultrafast universal RNA-seq aligner. Bioinformatics (Oxford, England) 29, 15–21. 10.1093/bioinformatics/bts635.

67. Li, B., and Dewey, C.N. (2011). RSEM: accurate transcript quantification from RNA-Seq data with or without a reference genome. Bmc Bioinformatics 12. Artn 323 10.1186/1471-2105-12-323.

68. Hanzelmann, S., Castelo, R., and Guinney, J. (2013). GSVA: gene set variation analysis for microarray and RNA-Seq data. Bmc Bioinformatics 14. Artn 7 10.1186/1471-2105-14-7.

69. Gu, Z.G., Eils, R., and Schlesner, M. (2016). Complex heatmaps reveal patterns and correlations in multidimensional genomic data. Bioinformatics 32, 2847–2849. 10.1093/bioinformatics/btw313.

70. Shen, S.H., Park, J.W., Lu, Z.X., Lin, L., Henry, M.D., Wu, Y.N., Zhou, Q., and Xing, Y. (2014). rMATS: Robust and flexible detection of differential alternative splicing from replicate RNA-Seq data. P Natl Acad Sci USA 111, E5593–E5601. 10.1073/pnas.1419161111.

71. Howe, K.L., Achuthan, P., Allen, J., Allen, J., Alvarez-Jarreta, J., Amode, M.R., Armean, I.M., Azov, A.G., Bennett, R., Bhai, J., et al. (2021). Ensembl 2021. Nucleic Acids Res 49, D884–D891. 10.1093/nar/gkaa942.

72. Stuart, T., Srivastava, A., Madad, S., Lareau, C.A., and Satija, R. (2021). Single-cell chromatin state analysis with Signac. Nat Methods 18, 1333-+. 10.1038/s41592-021-01282-5.

73. Zhang, Y., Liu, T., Meyer, C.A., Eeckhoute, J., Johnson, D.S., Bernstein, B.E., Nussbaum, C., Myers, R.M., Brown, M., Li, W., and Liu, X.S. (2008). Model-based Analysis of ChIP-Seq (MACS). Genome Biol 9. ARTN R137 10.1186/gb-2008-9-9-r137.

74. Cao, J.Y., Spielmann, M., Qiu, X.J., Huang, X.F., Ibrahim, D.M., Hill, A.J., Zhang, F., Mundlos, S., Christiansen, L., Steemers, F.J., et al. (2019). The single-cell transcriptional landscape of mammalian organogenesis. Nature 566, 496-+. 10.1038/s41586-019-0969-x.

75. Qiu, X.J., Mao, Q., Tang, Y., Wang, L., Chawla, R., Pliner, H.A., and Trapnell, C. (2017). Reversed graph embedding resolves complex single-cell trajectories. Nat Methods 14, 979-+. 10.1038/Nmeth.4402.

76. Meers, M.P., Tenenbaum, D., and Henikoff, S. (2019). Peak calling by Sparse Enrichment Analysis for CUT&RUN chromatin profiling. Epigenet Chromatin 12. ARTN 42 10.1186/s13072-019-0287-4.

77. Li, H.-D., Lin, C.-X., and Zheng, J. (2022). GTFtools: a software package for analyzing various features of gene models. Bioinformatics (Oxford, England) 38, 4806–4808. 10.1093/bioinformatics/btac561.

78. Loven, J., Hoke, H.A., Lin, C.Y., Lau, A., Orlando, D.A., Vakoc, C.R., Bradner, J.E., Lee, T.I., and Young, R.A. (2013). Selective Inhibition of Tumor Oncogenes by Disruption of Super-Enhancers. Cell 153, 320–334. 10.1016/j.cell.2013.03.036.

79. Ross-Innes, C.S., Stark, R., Teschendorff, A.E., Holmes, K.A., Ali, H.R., Dunning, M.J., Brown, G.D., Gojis, O., Ellis, I.O., Green, A.R., et al. (2012). Differential oestrogen receptor binding is associated with clinical outcome in breast cancer. Nature 481, 389–U177. 10.1038/nature10730.

80. Yu, G.C., Wang, L.G., and He, Q.Y. (2015). ChIPseeker: an R/Bioconductor package for ChIP peak annotation, comparison and visualization. Bioinformatics 31, 2382–2383. 10.1093/bioinformatics/btv145.

81. Subramanian, A., Tamayo, P., Mootha, V.K., Mukherjee, S., Ebert, B.L., Gillette, M.A., Paulovich, A., Pomeroy, S.L., Golub, T.R., Lander, E.S., and Mesirov, J.P. (2005). Gene set enrichment analysis: A knowledge-based approach for interpreting genome-wide expression profiles. P Natl Acad Sci USA 102, 15545–15550. 10.1073/pnas.0506580102.

