## Supplementary Materials for "Defined cellular reprogramming of androgen receptor-active prostate cancer to neuroendocrine prostate cancer"

**SUPPLEMENTARY FIGURE LEGENDS**

**FigureS1.** **Sustained reprogramming of ARPC to NEPC and functional bypass of AR signaling.** (**A**) Photomicrographs of H&E- and IHC-stained tissue sections of C4-2B cell line xenograft tumors established subcutaneously in non-castrate adult male NGS mice. (**B**) Photomicrographs of clonal C4-2B ARE-FKBP-Casp8 cells transduced with either GFP control or the candidate factor LV pool and subjected to no treatment or treatment with R1881 and AP20187 for five days. Scale bar = 100 μm.

**FigureS2. ASCL1 drives NEtD and suppresses AR expression in LNCaP cells. (A)** Immunoblot analysis of leave-one-out conditions in reprogramming studies using the C4-2B cell line. Lysates were collected on day 14. **(B)** Plot of the distribution of barcodes identified by single-cell DNA amplicon sequencing per C4-2B cell 72 hours post transduction with the PRNBSA lentiviral pool. A total of 3,870 cells were analyzed. **(C)** Relative cell viability over time determined by CellTiter-Glo assay of LNCaP cells reprogrammed with various factor combinations. * denotes p <0.05.

**FigureS3. NEtD of MDA PCa 2b cells and evaluation of the competence of NE-associated transcription factors in driving NE reprogramming.** (**A**) Immunoblot analysis of MDA PCa 2b cell line conditions subjected to the NE transdifferentiation assay. (**B**) Immunoblot analysis of reprogrammed C4-2B cell line conditions in which NE-associated transcription factors were added to PRNB (dominant-negative TP53 H175R, shRB1, MYCN, and BCL2) and SRRM4 to evaluate their effects on markers of the AR and NE programs. (C) Heatmap of RNA-seq gene expression from reprogrammed MDA PCa 2b cell line conditions showing genes associated with the AR program and NE programs (NEURO I and NEURO II). UQ: upper quartile normalization. (**D**) Partial least squares-discriminant analysis (PLS-DA) plot based on RNA-seq gene expression of reprogrammed MDA PCa 2b cell line conditions (black shapes) projected onto human ARPC (red dots) and NEPC (blue dots) samples from Beltran et al., 2016. Ellipses represent 95% confidence level for multivariate t-distributions defined by ARPC (red) and NEPC (blue) data.

**FigureS4. SRRM4 expression leads to alternative splicing of REST in C4-2B reprogrammed cells.** (**A**) Sashimi plots of *REST* RNA-seq reads from C4-2B cells modified with PRNB and SRRM4 (PRNBS), PRNBS and NeuroD1 (PRNBSN), or PRNBS and ASCL1 (PRNBSA). Conditions with SRRM4 demonstrate inclusion of a neural exon leading to the REST-4 isoform. (**B**) PLS-DA plot based on alternative splicing analysis of C4-2B cells line conditions projected onto human ARPC (red dots) and NEPC (blue dots) samples from Beltran et al., 2016. Ellipses represent 95% confidence of t-distribution defined by ARPC (red) and NEPC (blue) data.

**FigureS5. ASCL1 or NeuroD1 but not SRRM4 in combination with PRNB is sufficient to reprogram ARPC to NEPC.** PCA analyses of bulk RNA-seq data from C4-2B cells reprogrammed with (**A**) PRNBS, (**B**) PRNBSA, and (**C**) PRNBSN over time. (**D**) PLS-DA plot based on RNA-seq gene expression of C4-2B cells reprogrammed with PRNBS over time (black shapes) projected onto human ARPC (red dots) and NEPC (blue dots) samples from Beltran et al., 2016. Ellipses represent 90% confidence for ARPC (red) and NEPC (blue).

**FigureS6. Temporal shift in global active enhancers in PRNBSA reprogrammed C4-2B cells.** (**A**) Heatmap showing dynamic changes in active enhancer regions defined by overlapping H3K4me1 and H3K27ac signals in C4-2B cells reprogrammed with PRNBSA over the 14-day reprogramming period. Plots showing Gene Ontology (GO) enrichments of MSigDB gene sets associated with (**B**) decreasing and (**C**) increasing active enhancer regions from D2 to D14 in C4-2B cells reprogrammed with PRNBSA. RLE: Relative Log Expression normalization. Genomics regions in (A) were obtained by setting a p-value cutoff of 1e-04 on Kendall correlation between day and peak activity. Red lines in (B) and (C) represent adjusted p-value = 0.05.

**FigureS7. Temporal shift in global active promoters in PRNBSA reprogrammed C4-2B cells.** (**A**) Heatmap showing significant changes in active promoter regions defined by overlapping H3K4me3 and H3K27ac signals in C4-2B cells reprogrammed with PRNBSA over the 14-day reprogramming period. RLE: Relative Log Expression normalization. Genomics regions were obtained by setting a p-value cutoff of 1e-02 on Kendall correlation between day and peak activity. (**B**) Plots showing GO enrichment of MSigDB gene sets associated with decreasing and increasing active promoter regions from D2 to D14 in in C4-2B cells reprogrammed with PRNBSA. Red lines represent adjusted p-value = 0.05.

**FigureS8. scRNA-seq profiles distinct cell states induced by NE reprogramming of C4-2B cells.** Monocle pseudotime trajectory plots of C4-2B cells reprogramming conditions. Cell populations were colored by (**A**) conditions, (**B**) pseudotime, (**C**), NE score, (**D**) *AR* expression, (**E**) *ASCL1* expression, and (**F**) *NEUROD1* expression.

**FigureS9. ASCL1 and NeuroD1 engage in self-regulation after NE reprogramming of PC.** (**A**) Experimental schema to evaluate exogenous and endogenous ASCL1 or NeuroD1 gene expression using differential usage analysis. (**B**) Integrated Genomics Viewer (IGV) tracks of RNA-seq reads from reprogrammed C4-2B cell line conditions mapping to *ASCL1* or the untranslated regions (UTRs) of *ASCL1*. The relative distribution of reads with the synonymous single-nucleotide variant NM_004316.4(ASCL1);c.627C>G encoded in the ASCL1 lentivirus is shown in the magnified image. (**C**) IGV tracks of RNA-seq reads from reprogrammed C4-2B cell line conditions mapping to *NEUROD1* or *NEUROD1* UTRs. The magnified tracks below show reads mapping to the *NEUROD1* UTRs associated with conditions where exogenous NeuroD1 was introduced.

**FigureS10. NE signature scores are associated with reduced MHC I antigen processing and presentation signature scores in PC.** (**A**) Correlations of gene expression and various MHC signature scores to NE signature scores in the ARPC and NEPC samples from Beltran et al., 2016. (**B**) Correlations of gene expression and various MHC signature scores to NE signature scores in the reprogrammed C4-2B cell lines. Correlation coefficients (R) and P-values (P) were derived from Pearson correlations.

**FigureS11. Downregulation of MHC class I antigen processing and presentation genes starts early during reprogramming of ARPC to NEPC.** Heatmap of RNA-seq gene expression data from C4-2B cells reprogrammed with PRNBSA or PRNBSN over the 14-day reprogramming period showing the NE signature and MHC class I pathway scores (top) and select MHC class I genes including *B2M* (bottom). UQ: upper quartile normalization.

**FigureS12. Increased NE signature are associated with downregulation of MHC class I antigen processing and presentation genes in MDA PCa cell lines. (A)** Heatmap of RNA-seq gene expression data from reprogrammed MDA PCa cell line conditions showing the NE signature and MHC class I pathway scores (top) and select MHC class I genes including B2M (bottom). UQ: upper quartile normalization. **(B)** Correlations of gene expression and various MHC signature scores to NE signature scores in the reprogrammed MDA PCa cell lines. Correlation coefficients (R) and P-values (P) were derived from Pearson correlations.

**SUPPLEMENTARY FIGURES**

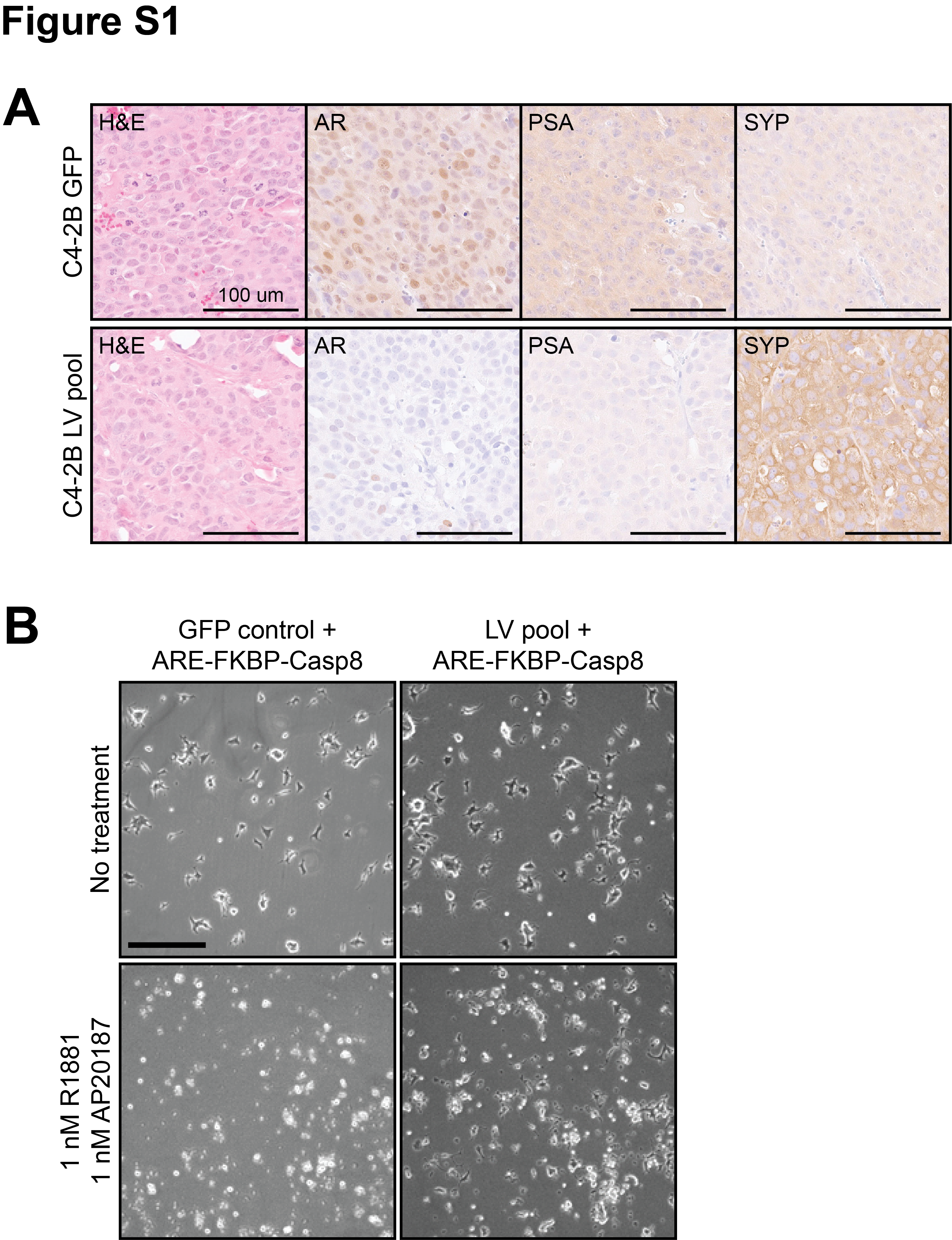

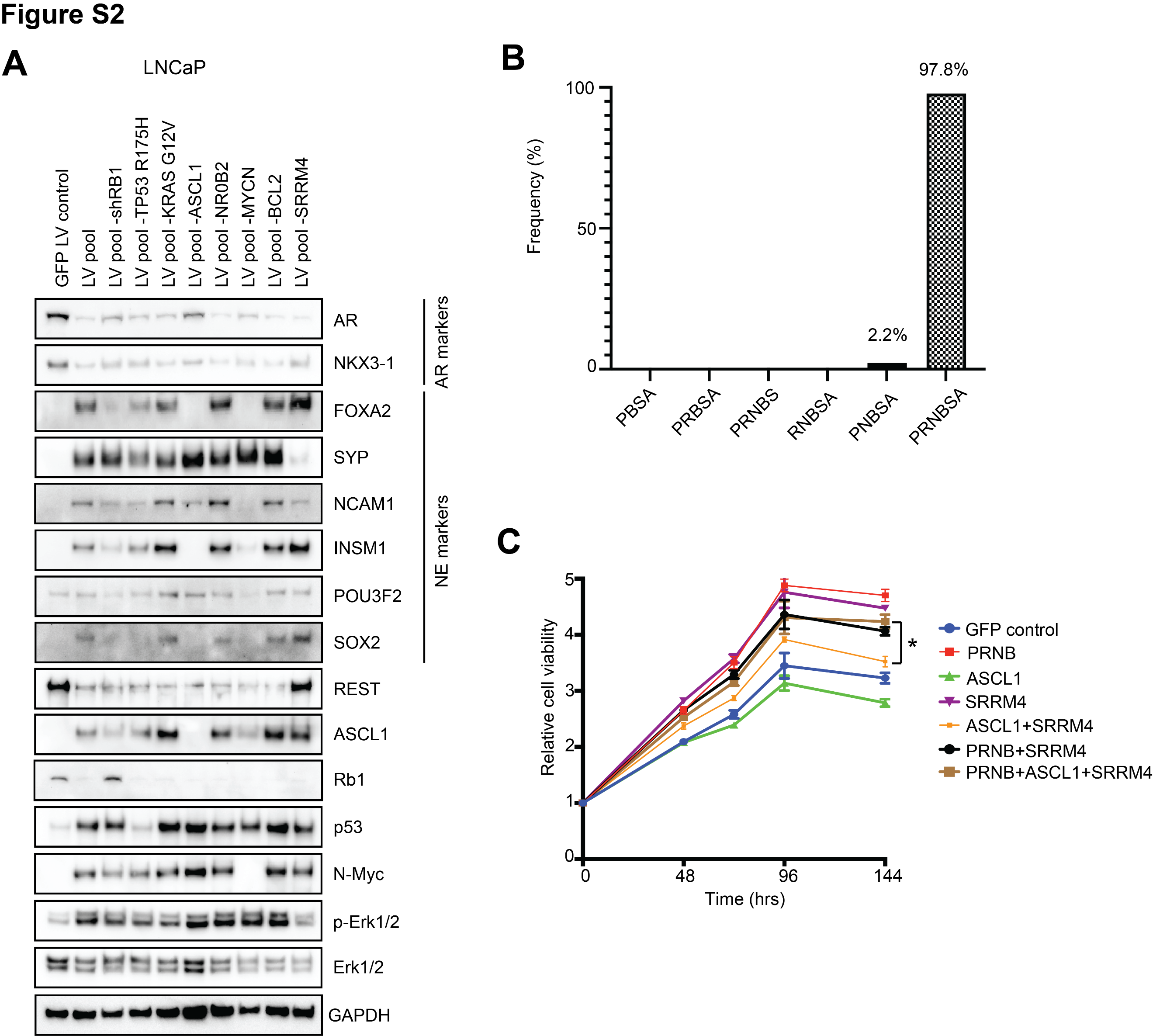

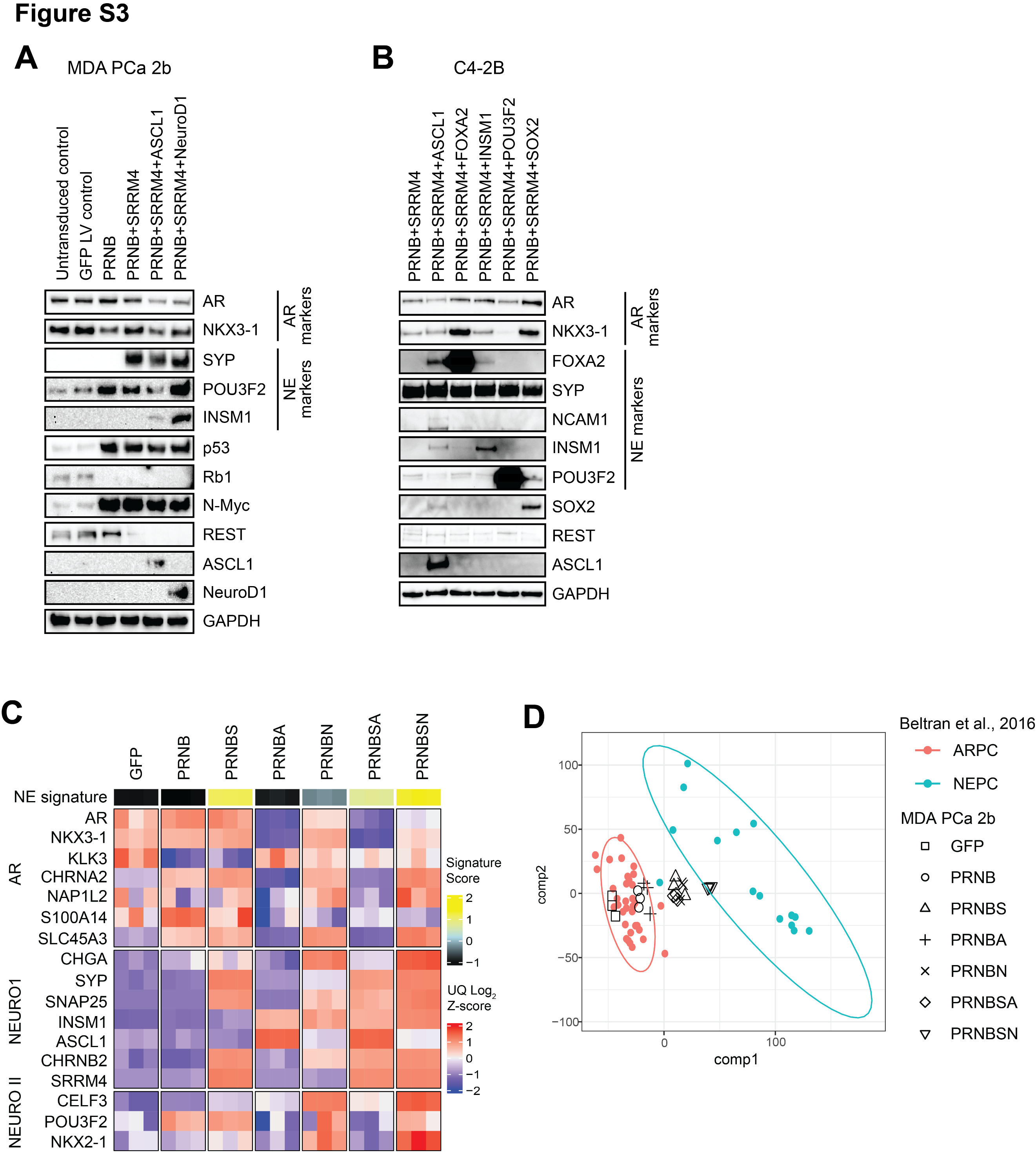

**­**
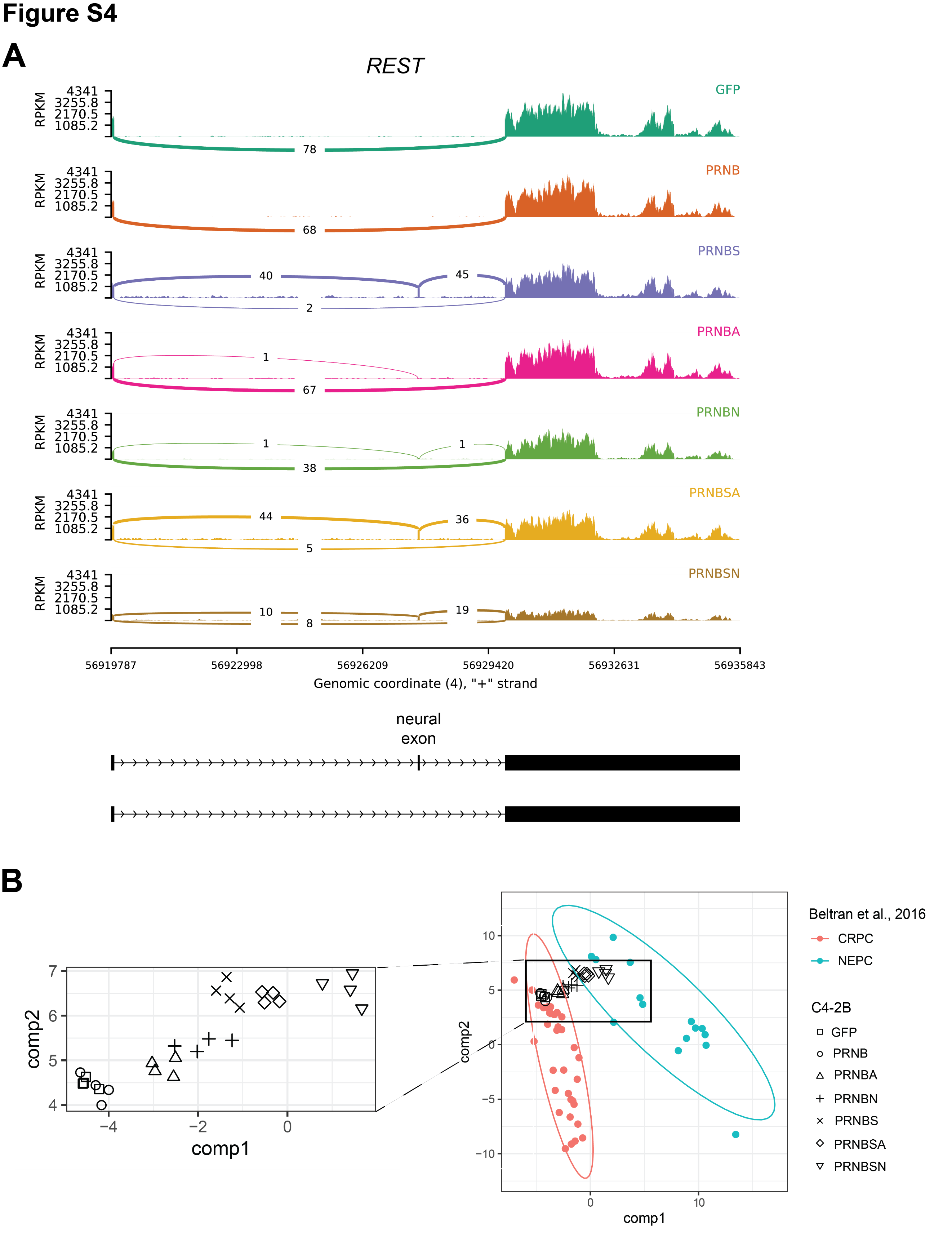

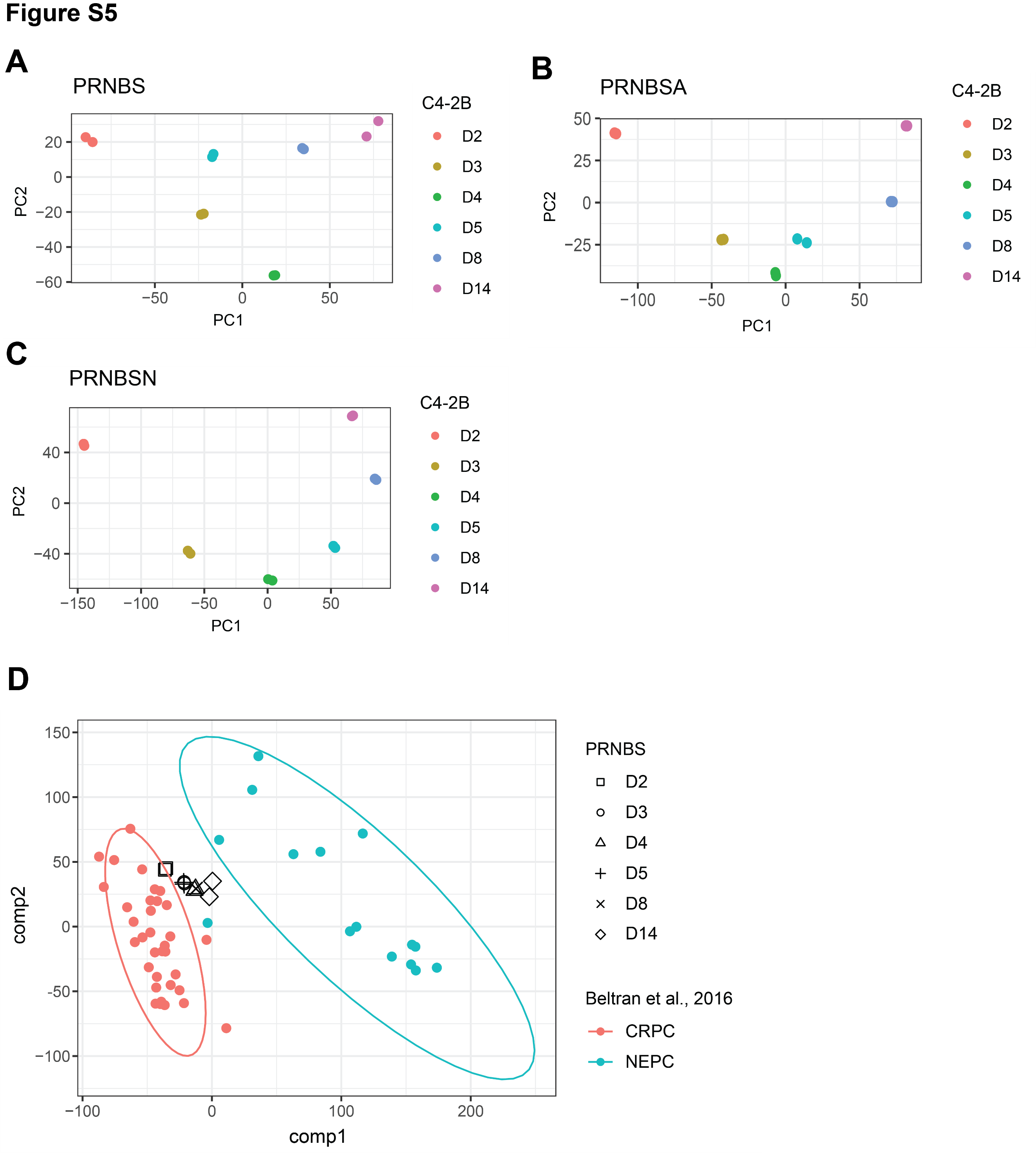

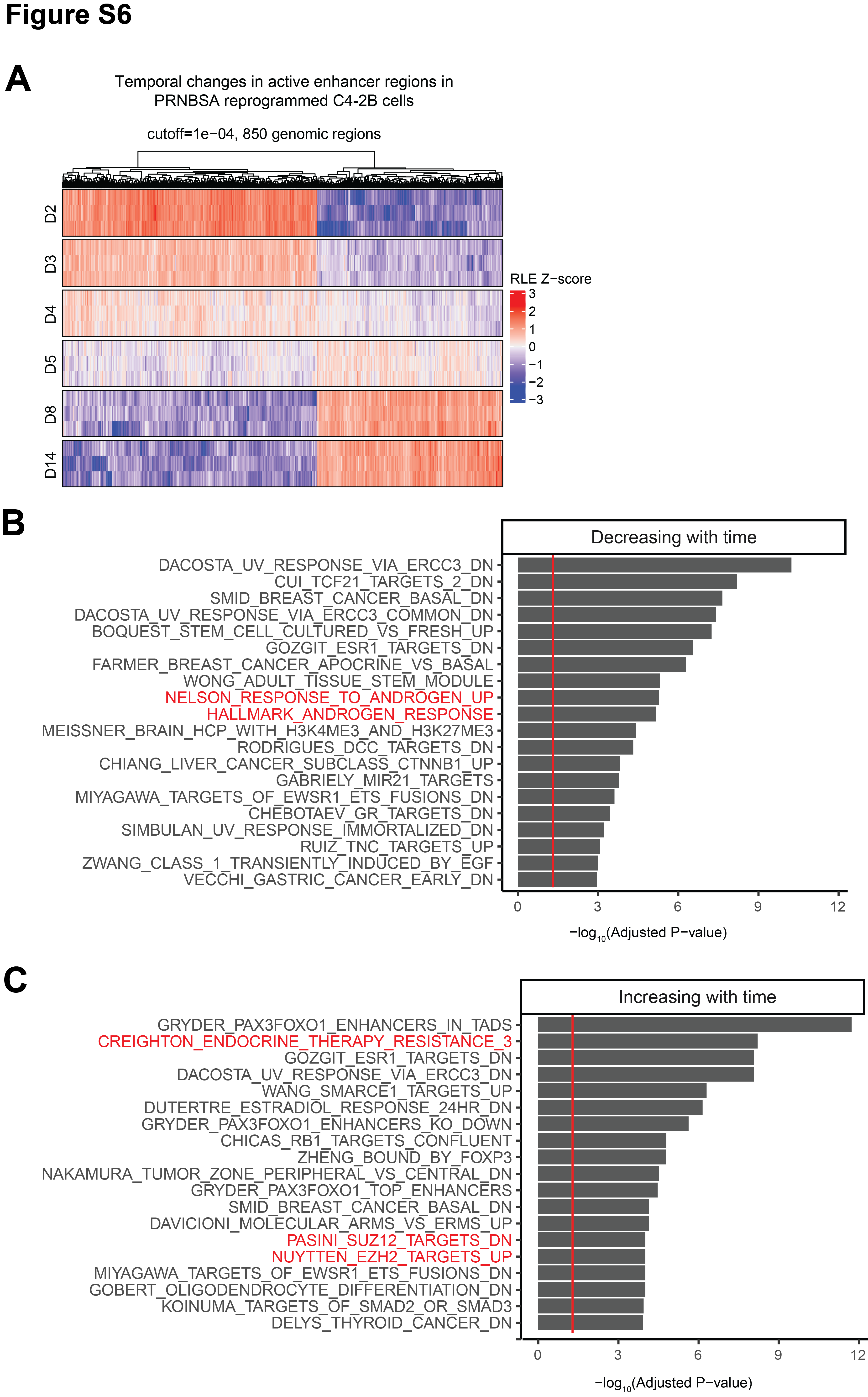

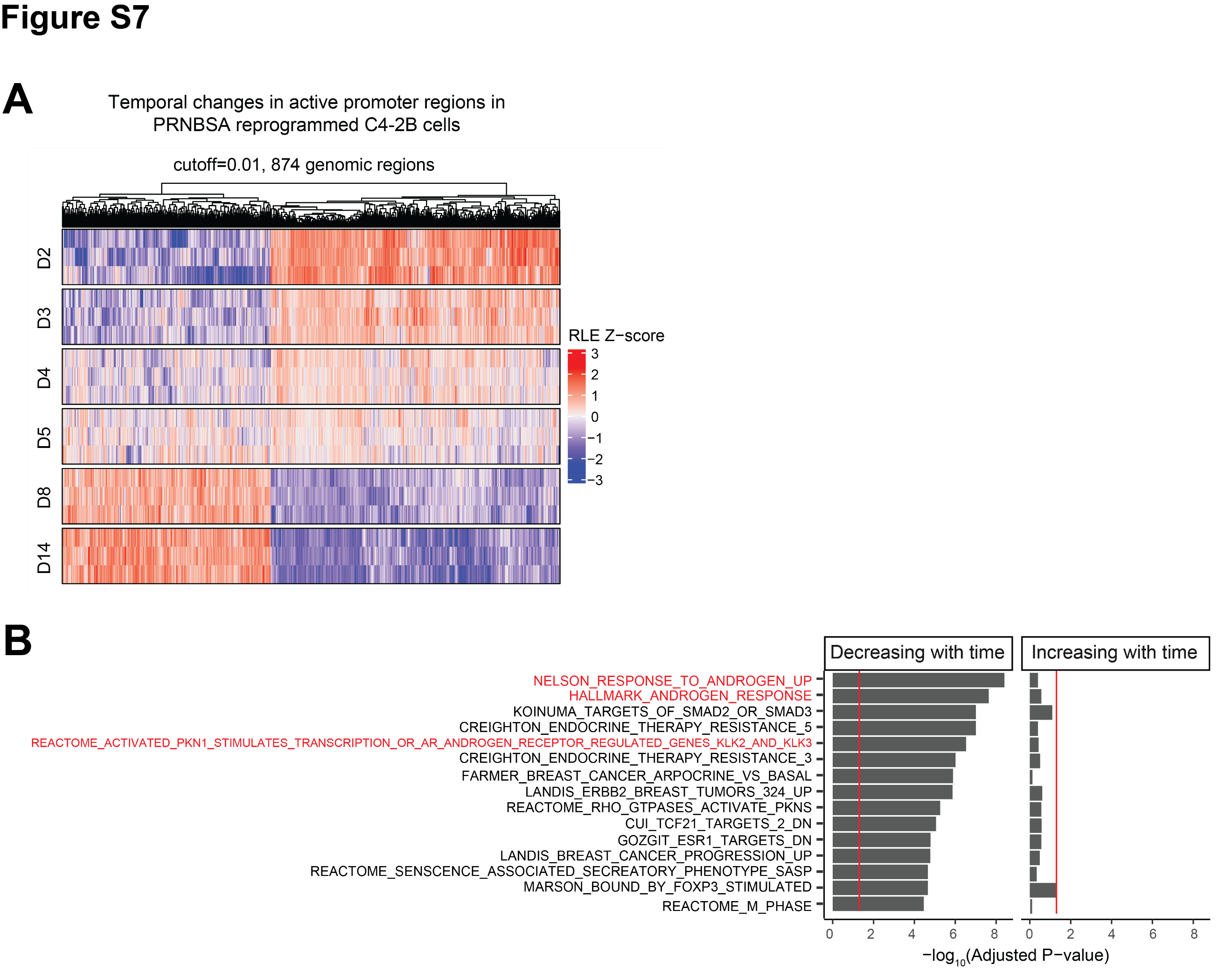

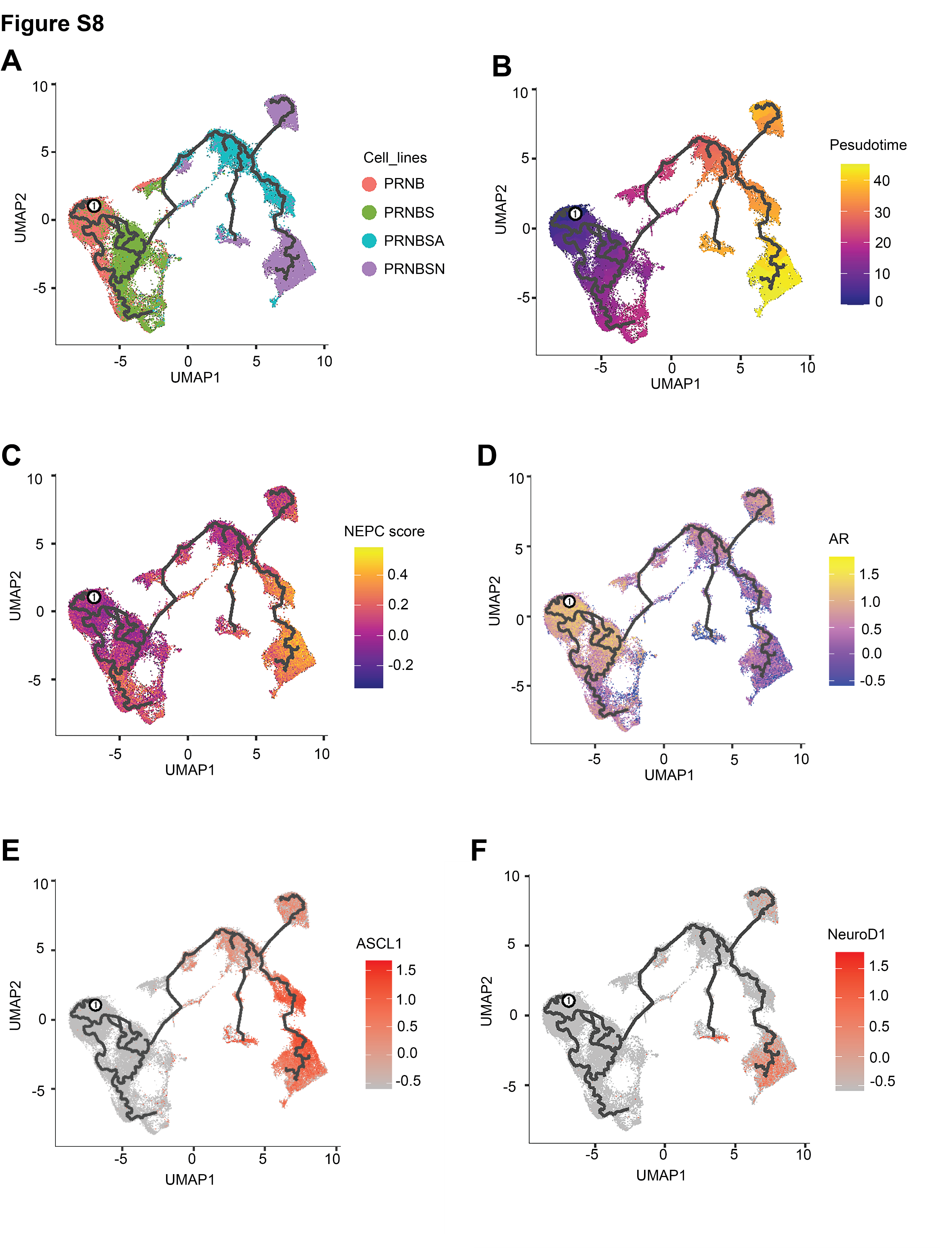

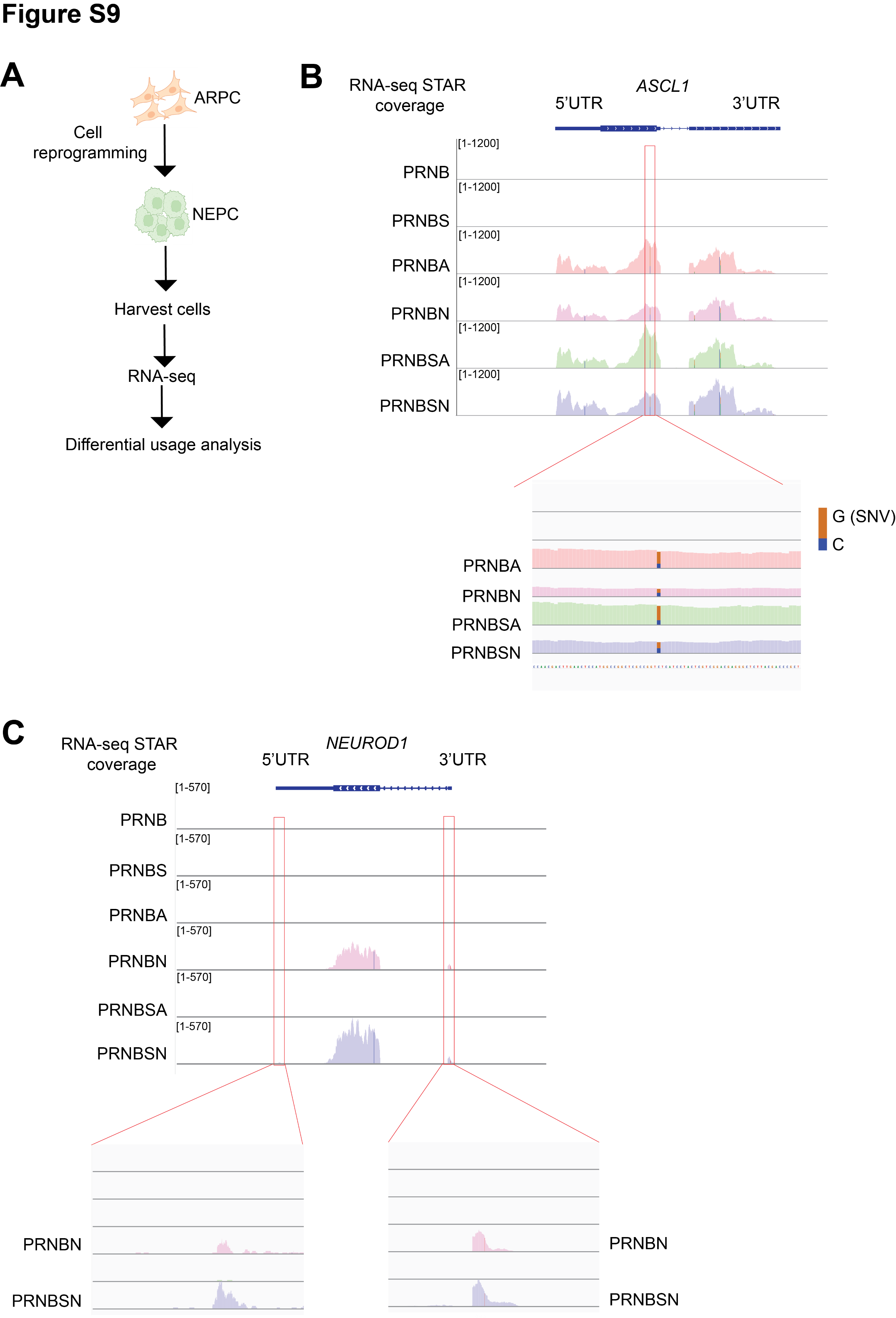

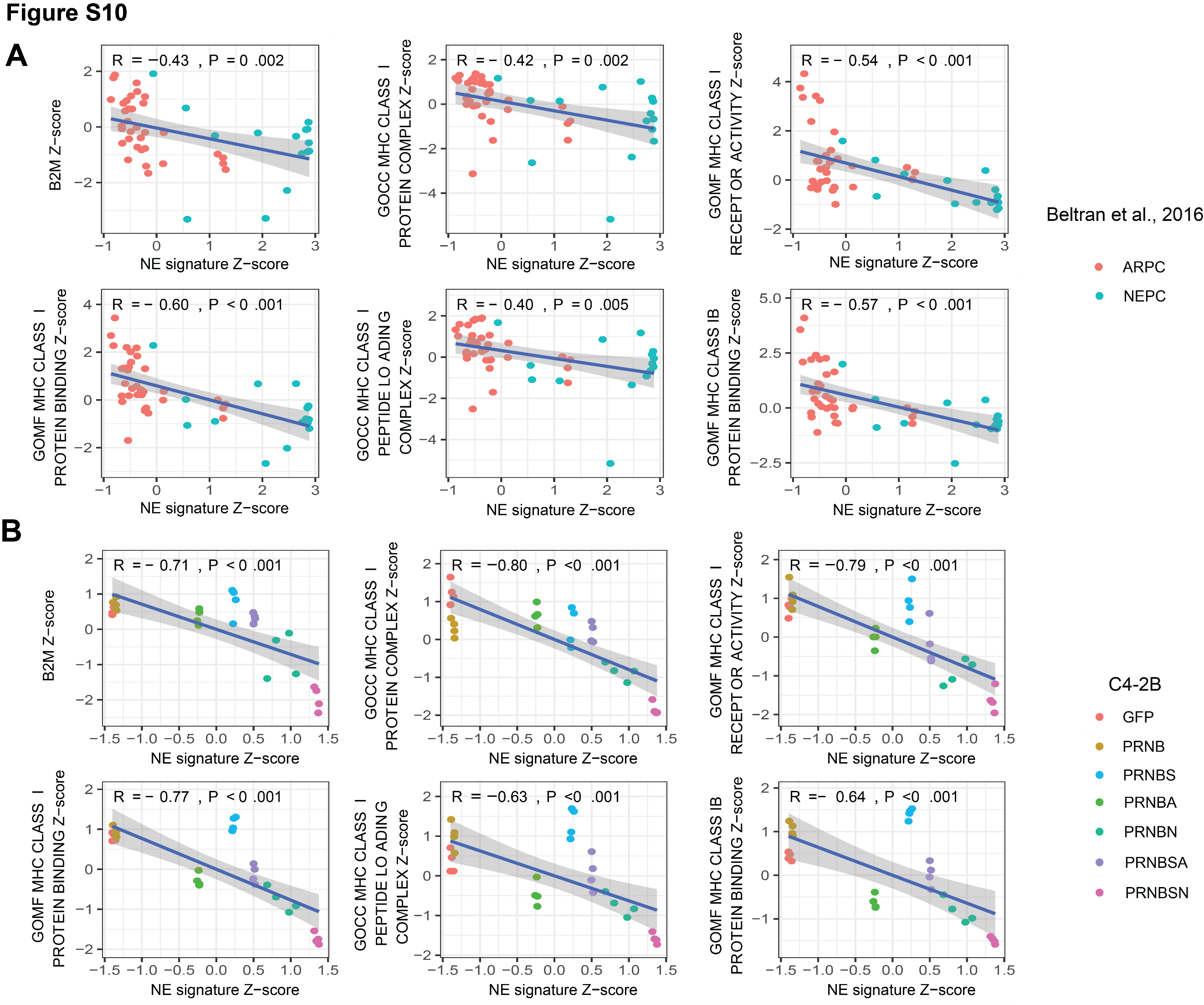

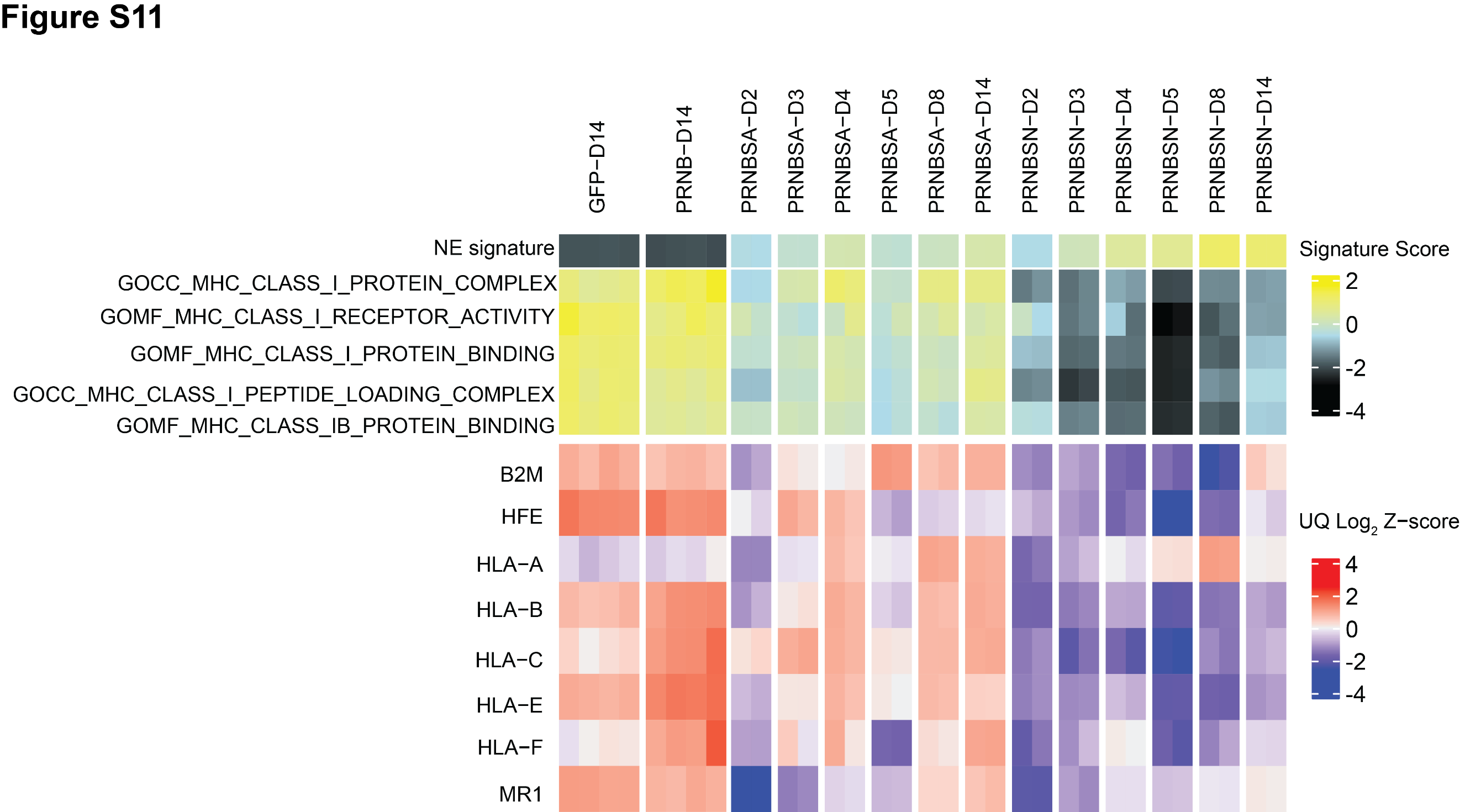

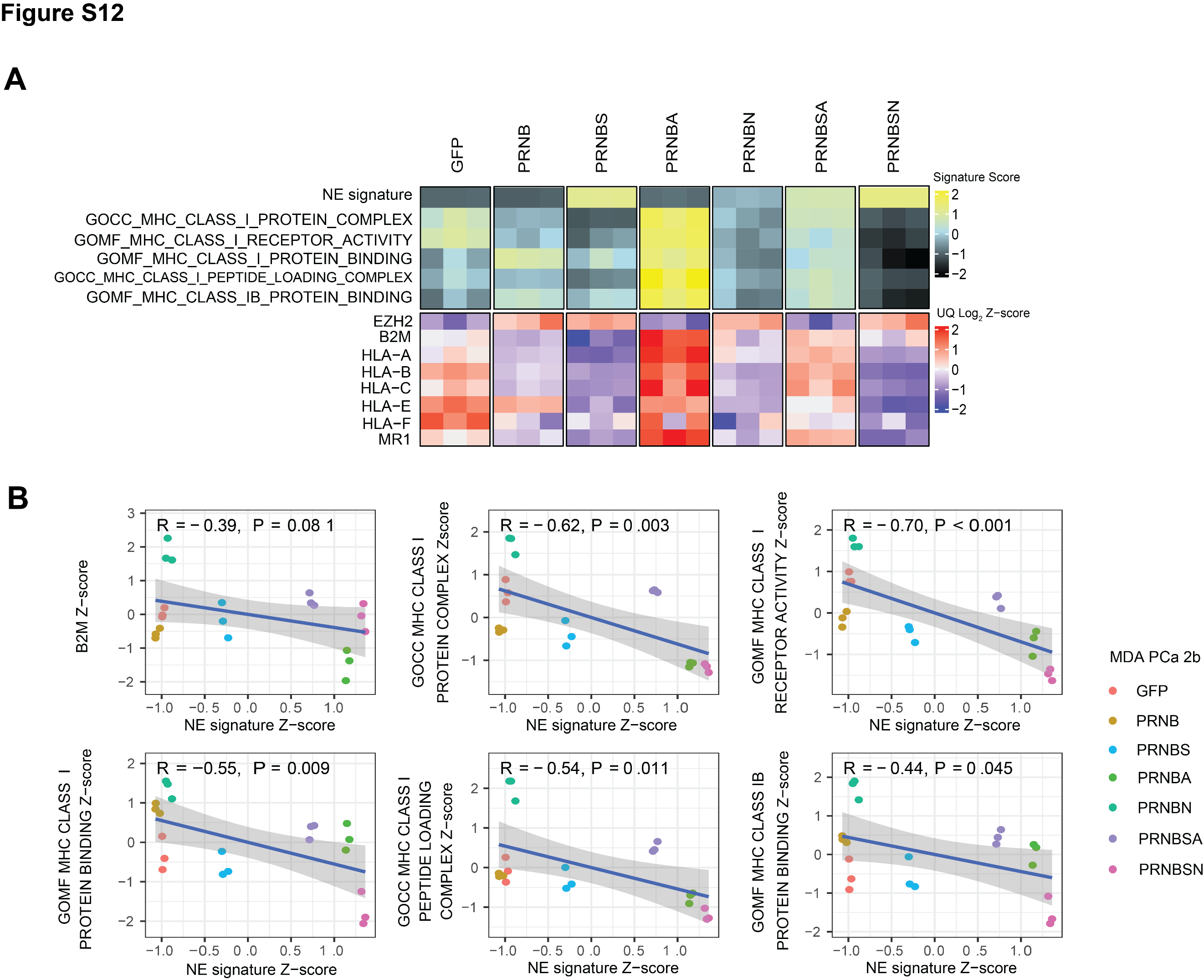

**Table S1**. **List of antibodies used for Western Blotting (WB), Immunohistochemistry (IHC), Immunocytochemistry (ICC), Immunofluorescence (IF), and Cleavage Under Targets and Release Using Nuclease (CUT&RUN).**

| **Antibodies** | **Application** | **Source** | **Catalog number** | **Dilution** | **Species** |
| --- | --- | --- | --- | --- | --- |
| AR | WB, ICC,IHC | Santa Cruz | sc-816 | 1:1000 | mouse |
| AR | IF | Cell Signaling | 5153T | 1:100 | rabbit |
| ASCL1 (24B72D11.1) | WB | BD Pharmingen | 556604 | 1:1000 | mouse |
| ASCL1 (24B72D11.1) | IF | BD Pharmingen | 556604 | 1:25 | mouse |
| N-MYC | WB | Santa Cruz | sc-53993 | 1:1000 | mouse |
| RB1 | WB | Santa Cruz | sc-74562 | 1:500 | mouse |
| P53 (D0-1) | WB | Santa Cruz | sc-126 | 1:500 | mouse |
| BCL2 (124) | WB | Cell Signaling | 15071S | 1:1000 | mouse |
| NeuroD1 (A-10) | WB | Santa Cruz | sc-46684 | 1:500 | mouse |
| NKX3-1 | WB | Cell Signaling | 83700S | 1:2000 | rabbit |
| INSM1 | WB | Santa Cruz | sc-271408 | 1:500 | mouse |
| NCAM | WB | Santa Cruz | sc-374289 | 1:500 | mouse |
| FOXA2 | WB | Santa Cruz | sc-377033 | 1:500 | mouse |
| SYP | WB, ICC, IHC | Santa Cruz | sc-17750 | 1:500 | mouse |
| PSA | IHC | Santa Cruz | sc-7306 | 1:500 | mouse |
| POU3F2 | WB | Cell Signaling | 12137 | 1:1000 | rabbit |
| REST | WB | Proteintech | 22242-1-AP | 1:250 | mouse |
| SOX2 (E-4) | WB | Santa Cruz | sc-365823 | 1:1000 | mouse |
| Erk1/2 | WB | Cell Signaling | 4695 | 1:1000 | rabbit |
| Phospho Erk1/2 | WB | Cell Signaling | 9101 | 1:1000 | rabbit |
| GAPDH-HRP | WB | GeneTex | GTX627408-01 | 1:10,000 | mouse |
| Goat anti-rabbit-HRP Conjugate | WB | BioRad | 1706515 | 1:10,000 | mouse |
| Goat anti-mouse-HRP conjugate | WB | BioRad | 1706516 | 1:10,000 | mouse |
| ASCL1 | CUT&RUN | Abcam | ab74065 | 1:50 | rabbit |
| H3K4me1 | CUT&RUN | Abcam | ab8895 | 1:100 | rabbit |
| H3K4me3 (C42D8) | CUT&RUN | Cell Signaling | 9751 | 1:100 | rabbit |
| H3K27ac | CUT&RUN | EMD Millipore | MABE647 | 1:50 | rabbit |
| Rabbit IgG | CUT&RUN | EpiCypher | 13-0042 | 1:10 | rabbit |
